# Analysis of Endothelial-to-Haematopoietic Transition at the Single Cell Level identifies Cell Cycle Regulation as a Driver of Differentiation

**DOI:** 10.1101/2020.04.03.023762

**Authors:** Giovanni Canu, Emmanouil Athanasiadis, Rodrigo A. Grandy, Jose Garcia-Bernardo, Paulina M. Strzelecka, Ludovic Vallier, Daniel Ortmann, Ana Cvejic

**Affiliations:** Wellcome Trust–Medical Research Council Cambridge Stem Cell Institute, Cambridge, UK; Department of Surgery, University of Cambridge, Cambridge, UK; Department of Haematology, University of Cambridge, Cambridge, UK; Wellcome Trust Sanger Institute, Wellcome Trust Genome Campus, Cambridge, UK; GSK, Medicines Research Centre, Gunnels Wood Road, Stevenage, Hertfordshire, SG1 2NY, UK

## Abstract

Haematopoietic stem cells (HSC) first arise during development in the aorta-gonad-mesonephros (AGM) region of the embryo from a population of haemogenic endothelial cells which undergo endothelial-to-haematopoietic transition (EHT). Despite the progress achieved in recent years, the molecular mechanisms driving EHT are still poorly understood, especially in human where the AGM region is not easily accessible. In this study, we took advantage of a human pluripotent stem cell (hPSC) differentiation system and single-cell transcriptomics to recapitulate EHT *in vitro* and uncover mechanisms by which the haemogenic endothelium generates early haematopoietic cells. We show that most of the endothelial cells reside in a quiescent state and progress to the haematopoietic fate within a defined time window, within which they need to re-enter into the cell cycle. If cell cycle is blocked, haemogenic endothelial cells lose their EHT potential and adopt a non-haemogenic identity. Furthermore, we demonstrated that CDK4/6 and CDK1 play a key role not only in the transition but also in allowing haematopoietic progenitors to establish their full differentiation potential. Therefore, we propose a direct link between the molecular machineries that control cell cycle progression and EHT.

## BACKGROUND

The first self-renewing haematopoietic stem cells (HSCs) are generated from the haemogenic endothelium, a specialised population of endothelial cells, located in the aorta-gonad-mesonephros (AGM) region (1, 2, 3). This process is known as endothelial-to-haematopoietic transition (EHT) and is characterised by the appearance of intra-aortic haematopoietic clusters (IAHCs). IAHCs are physically associated with the haemogenic endothelium which is lining the ventral wall of the dorsal aorta in human (4, 5). One of the first events that precedes EHT is the expression of RUNX1 in a subset of endothelial cells. Thus, RUNX1 expression marks the haemogenic endothelium where IAHCs will subsequently emerge (6). It has been shown that RUNX1 activates the haematopoietic programme and at the same time mediates the upregulation of transcription factors (*e.g.* GFI1 and GFI1B) which in turn repress endothelial genes (7). This dual role of RUNX1 possibly depends on its crosstalk with other key regulators of haematopoiesis such as TAL1 and GATA2 (8, 9). In addition to the AGM, other secondary sites have been reported to produce HSCs from haemogenic endothelial cells through EHT later on during development, such as placenta, vitelline/umbilical arteries, and embryonic head (5, 10, 11, 12, 13, 14). These first HSCs migrate to the foetal liver where their number dramatically increases, both as a consequence of proliferation and due to the contribution of secondary haematopoietic sites (5, 14). Despite its importance, the mechanisms controlling EHT remain to be fully uncovered, especially in human where these developmental stages are difficult to access for obvious ethical reasons. To bypass these limitations, several groups have developed *in vitro* methods that recapitulate production of haematopoietic cells through the generation of an intermediate endothelial state (15, 16, 17, 18, 19, 20, 21).

Here we took advantage of human pluripotent stem cells (hPSCs) to model haematopoietic development *in vitro* and used single-cell transcriptomics to dissect this process. We show that distinct populations are generated during EHT, including a population of haematopoietic progenitor cells that have multilineage differentiation potential. Furthermore, we demonstrated a tight link between cell cycle progression and EHT. Indeed, endothelial cells are quiescent and re-enter cell cycle to differentiate into haematopoietic progenitor cells. Inhibition of the cell cycle blocks EHT and causes endothelial cells to lose haemogenic potential. Finally, we demonstrated that cell cycle regulators such as CDK4/6 and CDK1 are not only essential for EHT but also control the capacity of nascent haematopoietic progenitors to differentiate. Together, our results uncover new mechanisms controlling the production of definitive haematopoietic cells which will be essential not only to understand blood cell development but also to improve protocols for generating these cells *in vitro*.

## RESULTS

hPSC differentiation provides an *in vitro* model of endothelial-to-haematopoietic transition.

In order to gain insight into mechanisms driving human definitive haematopoiesis, we utilised a system for the differentiation of hPSCs (**Fig. 1a**) (22, 23). This *in vitro* system recapitulates a natural path of development that leads to the production of an intermediate population of endothelial cells with haemogenic potential. Between EHT Day 3 (D3) and EHT Day 5 (D5) these endothelial cells generate round clusters that progressively increase in size and release single haematopoietic cells in the culture medium (**Fig. 1b**). Importantly, these haematopoietic cells can further differentiate into myeloid cells, foetal γ-globin-producing erythroid cells (Fig. **1c**, **Supplementary Fig. 1a, b**) as well as T lymphocytes (23, 24, 25). Transcriptionally, the process is marked by the gradual downregulation of endothelial markers (*e.g. CDH5, VWF*) and concomitant upregulation of blood markers (*e.g. SPI1, KLF1*), with key haematopoietic stem/progenitor cells (HSPCs) genes peaking at EHT D3 (*e.g. RUNX1*, *MYB* and *GATA2*) (**Supplementary Fig. 1c**, **Fig. 1d, e**). Taken together, our data show that differentiation of hPSCs morphologically resembles the generation of IAHCs (1, 2, 5, 25) and that the production of haematopoietic progenitors *in vitro* culminates around 3-5 days of EHT culture (**Supplementary Fig. 1c)**. For the aforementioned reasons, we selected this interval for our subsequent analyses.

**Figure 1.**
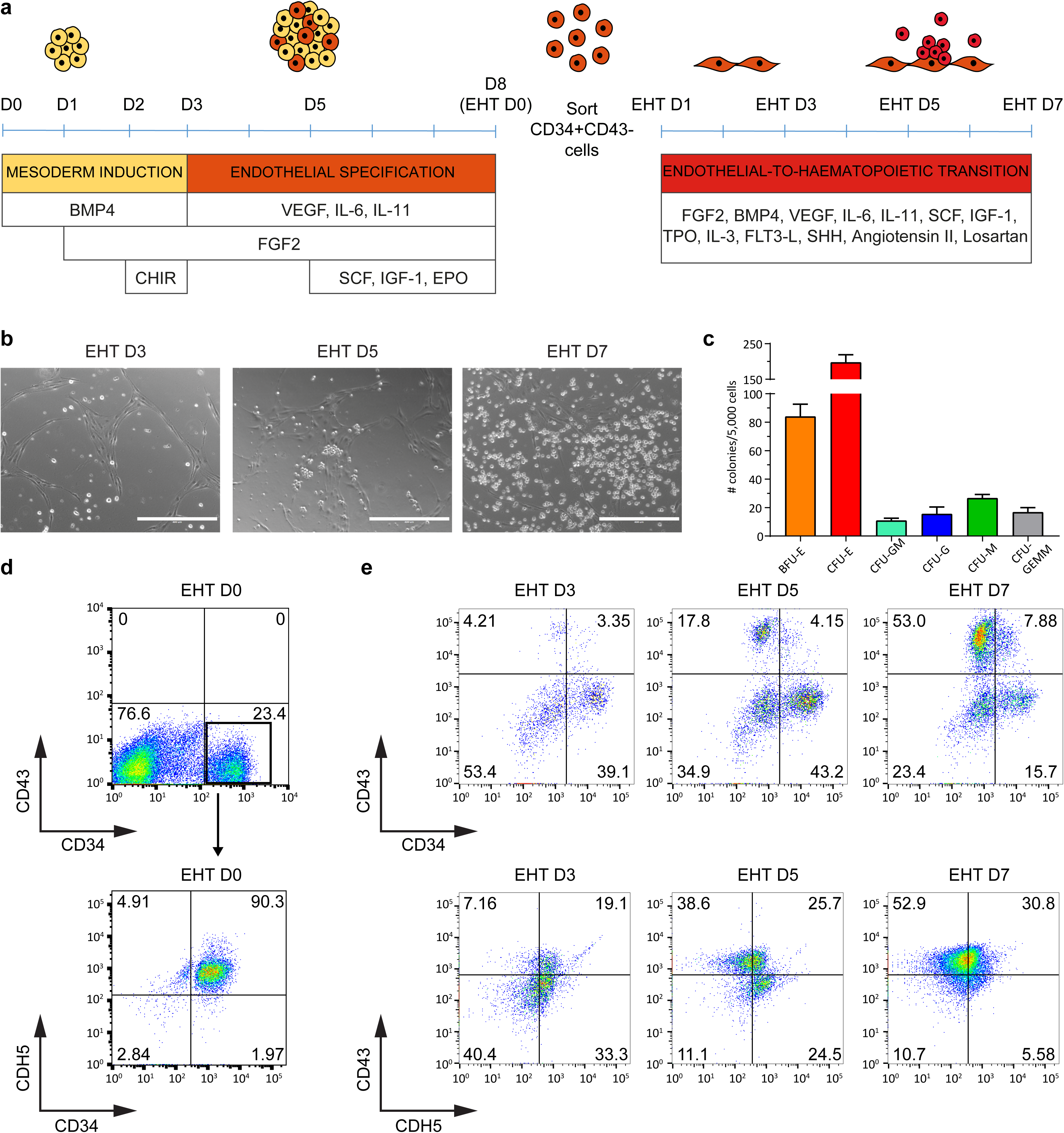
Culture system for modelling EHT *in vitro*. (A) Schematic of the differentiation protocol used to model EHT with hPSCs. D: day. **(B)** Representative images of cells progressing through EHT. Scale bar is 400 μm. **(C)** Quantification of colonies generated in a CFU assay using cells collected at EHT D5. Data represent means ±SEM of 3 independent experiments. BFU-E: burst forming unit-erythroid, CFU-E: colony forming unit-erythroid, CFU-GM: colony forming unit-granulocyte-macrophage, CFU-G: colony forming unit-granulocyte, CFU-M: colony forming unit-macrophage, CFU-GEMM: colony forming unit-granulocyte-erythrocyte-macrophage-megakaryocyte. **(D and E)** Flow cytometry analysis of (D) cells at D8/EHT D0 and (E) during EHT culture at multiple time points. Results are representative of at least 3 independent experiments.

### Single-cell analysis revealed four distinct populations during *in vitro* EHT

EHT is a dynamic process with different cell types and states coexisting, thereby rendering bulk analyses difficult to interpret. To overcome this limitation and resolve the inherent heterogeneity of cell types generated during *in vitro* differentiation, we took advantage of single-cell RNA sequencing (scRNA-seq). In brief, cells were collected at EHT D3 and D5, processed using the Chromium 10X Genomics system and analysed following the Seurat workflow (26). Following quality control, 3,877 cells from EHT D3 and 2,152 from EHT D5 were used in downstream analyses. In both datasets we identified four distinct populations (**Fig. 2a, b**) that were annotated based on marker genes and the presence of known lineage-specific signature genes (**Fig. 2c**, **Supplementary Fig. 2a**). Specifically, we identified a population of endothelial cells, positive for *CDH5* and *PECAM1*, mesenchymal cells, positive for *CDH2*, *KRT8* and *TAGLN,* and two clusters of haematopoietic cells: haematopoietic progenitor cells (HPCs) and erythroid cells. Cells in the HPC cluster displayed expression of stem cell genes such as *RUNX1* and *GATA2*, as well as low expression of early erythroid and myeloid genes, *e.g.*, *GATA1* and *CD33*. Cells in the erythroid cluster expressed erythroid lineage genes, *e.g.*, *KLF1*, *ALAS2* and *GYPA* (**Fig. 2c**, **Supplementary Fig. 2a)** and had immature cell morphology as shown by cytospin staining (**Supplementary Fig. 2b**). This cluster was therefore annotated as early erythroid progenitors.

**Figure 2.**
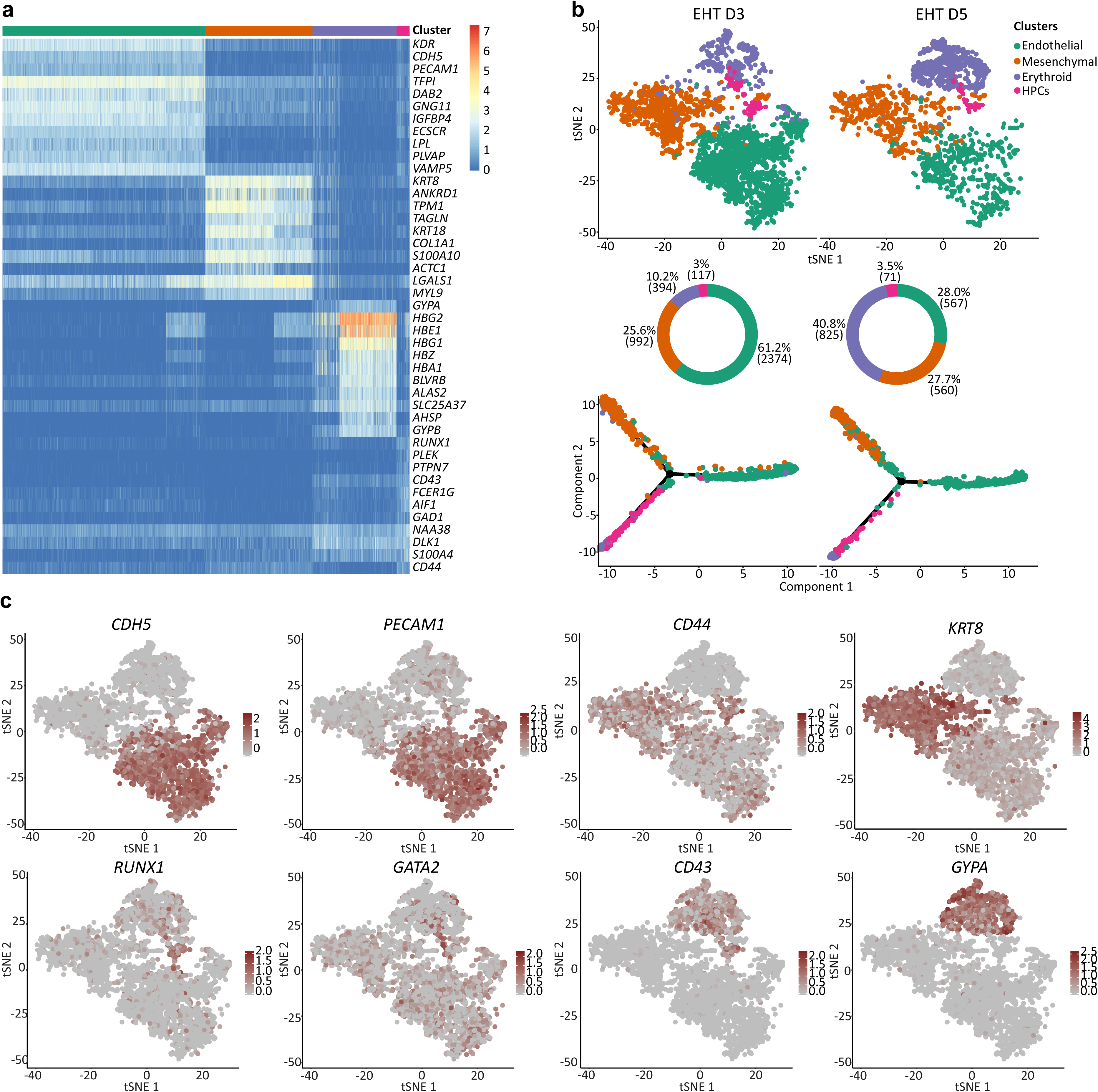
scRNA-seq reveals distinct cell types generated during hPSCs differentiation. (A) Heatmap of the marker genes for each cluster. For this analysis, data from EHT D3 and EHT D5 were merged. **(B)** tSNE plots showing comparison of samples collected at EHT D3 and EHT D5. Four populations were identified based on transcriptional identity. Pie charts are showing the percentage and absolute number (in brackets) of cells in each cluster. Transcriptional data was used to order cells along differentiation trajectories, from endothelial cells (right) to the haematopoietic and mesenchymal branches (left). **(C)** tSNE plots of merged data from EHT D3 and EHT D5. Expression of relevant lineage markers is shown.

To reconstruct the differentiation trajectory of haematopoietic cells during EHT, we compared cells from EHT D3 and EHT D5 (**Fig. 2b**). Our analyses revealed a marked reduction of endothelial cells and an increase of erythroid progenitors. Conversely, HPCs and mesenchymal cells only showed a minor increase. Pseudotime differentiation trajectories suggested that the haematopoietic and mesenchymal compartments both derived from a common endothelial population. Along this differentiation trajectory, endothelial cells gradually downregulated endothelial specific genes and concomitantly upregulated either mesenchymal or haematopoietic markers (**Supplementary Fig. 3a**). To validate the pseudotime differentiation trajectory, we sequenced further 1,764 CD34+/CD43-cells sorted at EHT D0 (**Supplementary Fig. 3b**). Our analysis demonstrated that most cells at EHT D0 were endothelial, as shown by the positive expression of multiple endothelial marker genes (*e.g. CDH5*, *PECAM1*, *KDR*, *THY1*). Importantly, we did not observe expression of haematopoietic lineage associated genes, demonstrating that haematopoietic cells are subsequently generated. Accordingly, cells collected at EHT D0 were not able to generate blood colonies in a CFU assay. In addition, none of the identified clusters had a clear mesenchymal transcriptional signature. However, a small population co-expressed endothelial (*e.g. KDR*, *THY1*) and mesenchymal markers (*e.g. COL1A1*, *CDH2*, *CDH11*, *KRT8*) at low level, suggesting that a fraction of cells displayed a mixed endothelial/mesenchymal identity at EHT D0 and represented an early stage of mesenchymal commitment.

### Identification and functional validation of multipotent HPCs

Based on our scRNA-seq data, we devised a sorting strategy to isolate each of the populations identified above. We predicted that combinations of three cell surface markers (*CDH5, CD43* and *CD44*) (**Fig. 2c**) was sufficient to allow sorting of the four populations. *CDH5* was mainly expressed in the endothelial cluster and to a lower extent in HPCs. The pan-haematopoietic *CD43* was expressed in HPCs and erythroid progenitors (**Fig. 2c**, **Supplementary Fig. 2a**). Both these genes are known to mark emerging HSPCs in the human dorsal aorta during development (32, 33). Finally, *CD44* appeared to be prevalently expressed in mesenchymal cells and HPCs (**Fig. 2c**). CD44 was previously shown to be expressed on IAHCs emerging during EHT in both human and mouse (34, 35, 36).

Immunostaining analyses confirmed that CDH5, CD43 and CD44 were co-expressed on the clusters of cells produced at EHT D5 (**Fig. 3a**). In addition, these cells expressed the master regulator of definitive haematopoiesis RUNX1 and were therefore likely to correspond to the HPC cluster identified in our single-cell analysis (**Supplementary Fig. 4a**). A lower expression of RUNX1 was also visible in some of the surrounding endothelial cells, possibly marking haemogenic endothelial cells in transition to the haematopoietic fate, consistent with what previously described *in vivo* (6).

**Figure 3.**
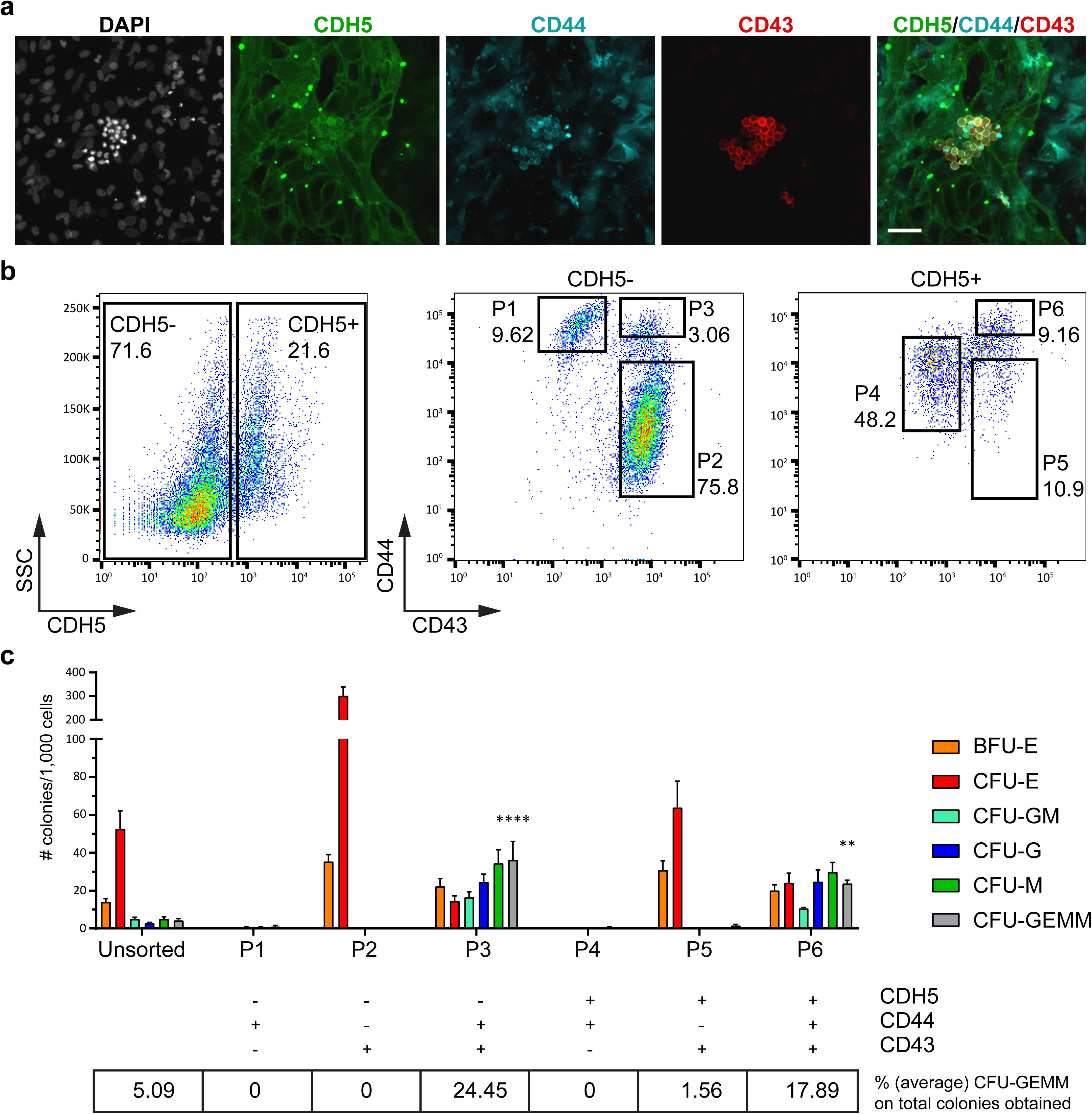
Sorting and functional validation of distinct populations. (A) Immunofluorescence staining showing expression of CDH5, CD44 and CD43 on haematopoietic clusters of cells at EHT D5. Scale bar is 50 μm. **(B)** Flow cytometry analysis showing the sorting strategy for six populations (P1-P6). Results are representative of at least 3 independent experiments, using 2 distinct hPSC lines. **(C)** CFU assay showing the differentiation potential of cells isolated from P1-P6. The ability to produce multilineage CFU-GEMM colonies is compared to the unsorted population. Data are mean ±SEM of 4 independent experiments, with statistical significance compared to control as *P<0.5, **P<0.01, ***P<0.001, ****P<0.0001 by one-way ANOVA. Degrees of freedom DFn=6, DFd=21, value F=15.21.

Next, using combinations of the three cell surface markers, we isolated six distinct populations (**Fig. 3b**) which were assessed for their capacity to generate blood cells in a CFU assay (**Fig. 3c**). Cells in P1 (CDH5-/CD44+/CD43-) and P4 (CDH5+/CD44+/CD43-), predicted to label mesenchymal and endothelial populations respectively, did not show haematopoietic potential in CFU assay. To further confirm their mesenchymal and endothelial identity, cells from P1 and P4 gates were re-plated and further cultured (**Supplementary Fig. 4 b-g**). After 10 days in culture, more than half of the cells from P4 were CD31+/CD90+, consistent with an endothelial identity. Importantly, these cells were able to produce new mesenchymal (CDH5-/CD44high) and haematopoietic (CD43+) cells, confirming their differentiation potential. In contrast, cells sorted and re-plated from P1 gate displayed morphology and markers (CD44+/Vimentin+) consistent with a mesenchymal identity. These mesenchymal cells showed no ability to differentiate into endothelial cells (*i.e.* all cells were CDH5-). Furthermore, cell from the P2 gate (CDH5-/CD44-/CD43+), predicted to label the erythroid population, and P5 (CDH5+/CD44-/CD43+), exclusively generated erythroid colonies in a CFU assay (**Fig. 3c**). In contrast, cells in P3 (CDH5-/CD44+/CD43+) and P6 (CDH5+/CD44+/CD43+), predicted to label multipotent HPCs, were able to generate an array of colonies (**Fig. 3c**) including multilineage CFU-GEMM colonies at a higher rate when compared to unsorted cells. In addition, we monitored the expression of CD34, CD43, CD44 and CDH5 by flow cytometry, from EHT D0 to EHT D3 (**Supplementary Fig. 5**). Our time course experiment showed that the described P1-P6 populations are generated over time starting from a population of CD34+/CD43-/CD44+low cells. Importantly, this initial population of cells at EHT D0 was devoid of any haematopoietic potential.

Although both CDH5+ (P6) and CDH5-(P3) cells displayed multilineage differentiation potential (as shown by CFU assays), immunofluorescence staining showed that the haematopoietic clusters contained CDH5+ but not CDH5-cells (**Fig. 3a**). CDH5-HPCs could only be detected when both the attached and floating fractions of cells were analysed by flow cytometry, indicating that they were floating in the culture medium and were most likely derived from CDH5+ HPCs. Thus, variable expression of CDH5 in HPCs might indicate different stages of their differentiation toward the haematopoietic lineage, with CDH5+ cells representing earlier progenitors and CDH5-HPCs being more mature progenitors that have detached from the endothelium. This is in agreement with previous studies showing that HSCs emerging in the AGM express CDH5, which is then downregulated upon HSCs maturation (33).

In summary, these results validate the proposed cell identities and confirm their differentiation trajectories, demonstrating that endothelial cells are able to undergo EHT and endothelial-to-mesenchymal transition (EndoMT) (27, 28, 29, 30, 31) to generate haematopoietic and mesenchymal populations, respectively. Additionally, we identified a sorting strategy to enrich for HPCs generated *in vitro* and confirmed their multilineage haematopoietic potential using differentiation assays.

### Cell cycle progression is required for specification of HPCs

During the course of the experiments described above, we observed that clusters of HPCs contain cells with a distinct nuclear morphology compared to the surrounding endothelial and mesenchymal cells. Indeed, a majority of these cells displayed nuclear condensation typical of mitotic or dividing cells (**Fig. 3a**) thereby suggesting that HPCs represent a highly proliferative population. To confirm this observation, we used our scRNA-seq data to infer the cell cycle state of the HPCs, endothelial, mesenchymal and early erythroid populations (**Fig. 4a**) (26). While endothelial cells were enriched in G1, mesenchymal cells and HPCs displayed a more active cell cycle. In contrast, erythroid progenitors appeared to be extremely proliferative with only 2.5% of the cells in G1 (**Fig. 4a**). In line with this observation, genes belonging to the Rb and CIP/KIP families such as *RB1*, *RBL2/p130* and *CDKN1C/p57*, which inhibit G1 progression and G1/S transition, were gradually downregulated during the transition of endothelial cells to HPCs and upon their further differentiation to erythroid cells (**Supplementary Fig. 6a**). Concomitantly, proliferation markers such as *MKI67* and *CCNB2* were upregulated. Interestingly, we observed differential expression of cyclin D genes that are necessary for the activation of CDK4/6 and progression through the G1 phase. Endothelial cells were enriched for *cyclin D1* and *D2*, while HPCs appeared to switch to *cyclin D2* and *D3*. The more committed erythroid cluster preferentially expressed *cyclin D3* (**Supplementary Fig. 6a**). This might suggest a divergent requirement for cyclin D genes during EHT, and would be in agreement with genetic studies in the mouse showing that the simultaneous absence of the three cyclin Ds lead to severe haematopoietic defects (37).

**Figure 4.**
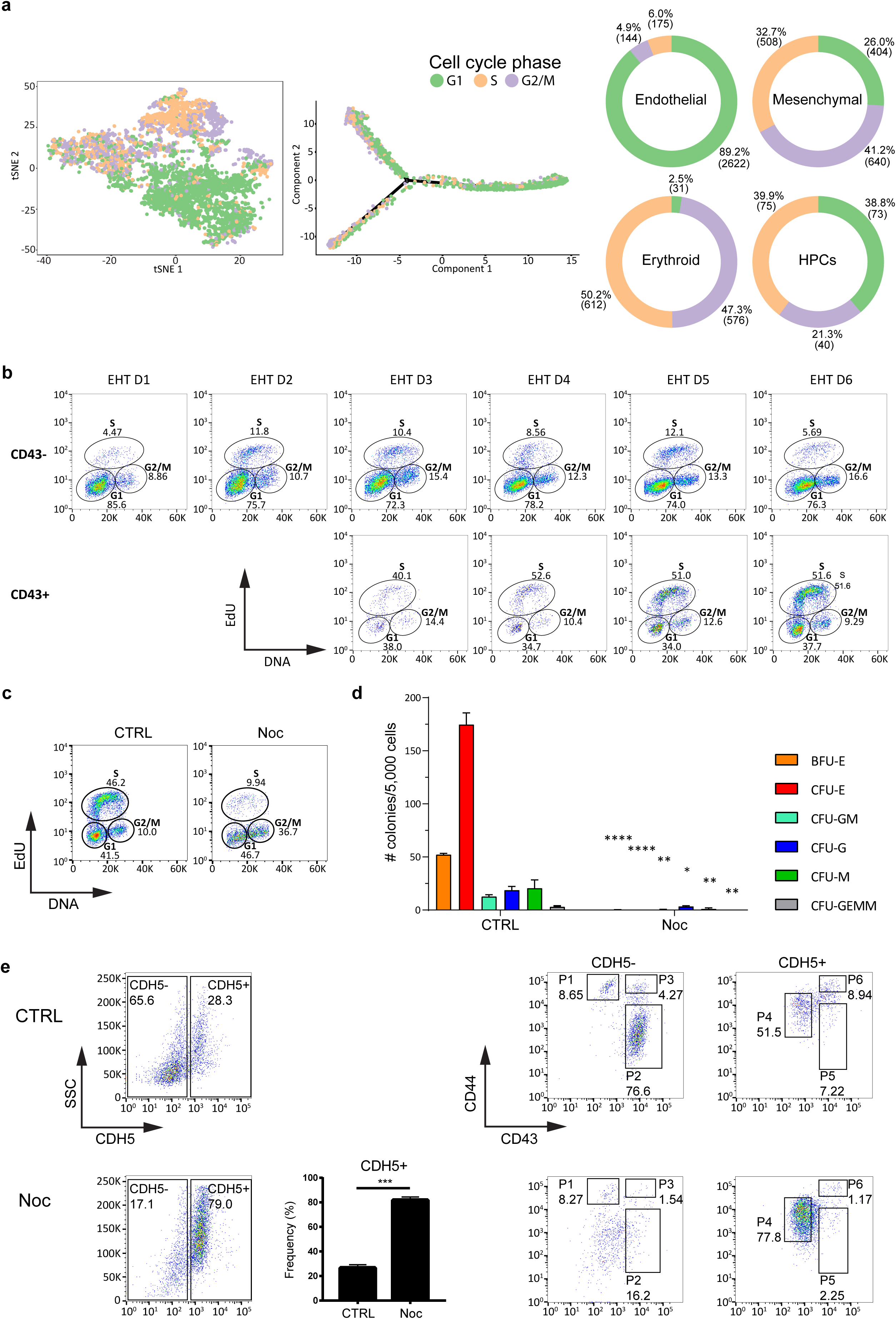
Cell cycle progression is necessary for EHT. (A) (Left panel) tSNE plot of the merged data from EHT D3 and EHT D5. Cells were coloured based on the cell cycle phase inferred from scRNA-seq data. (Middle panel) Cell cycle phase along differentiation trajectory, from endothelial cells (right) to haematopoietic and mesenchymal branches (left). (Right panel) Pie charts showing the percentage of cells in G1, G2/M and S phase in each of the clusters. **(B)** EdU staining and flow cytometry were used to monitor the cell cycle profile of CD43- and CD43+ cells during differentiation. Results are representative of 3 independent experiments. **(C – E)** At EHT D3 media was supplemented either with 0.1% DMSO (CTRL) or with 0.1 μg/mL Nocodazole (Noc). After 48h, at EHT D5 cells were either (C) processed for EdU staining and flow cytometry to monitor their cell cycle profile, (D) washed to remove Nocodazole and further cultured in a CFU assay to reveal their differentiation potential, or (E) stained for flow cytometry to monitor the distinct populations previously identified (P1: mesenchymal cells, P4: endothelial cells, P2 and P5: erythroid progenitors, P3 and P6: HPCs). Results are representative of 3 independent experiments. Data in (D) and bar plot in (E) are mean ±SEM of 3 independent experiments, with statistical significance compared to control as *P<0.5, **P<0.01, ***P<0.001, ****P<0.0001 by two-tailed unpaired t-test, and representative of 2 distinct hPSC lines. Degrees of freedom=4, t-values for the distinct colonies in (D) are BFU-E=39.62, CFU-E=15.74, CFU-GM=6.871, CFU-G=4.271, CFU-M=4.867, CFU-GEMM=5.196, t-value in (E) =15.05.

To experimentally validate these observations, we assessed the cell cycle profile of CD43+ (haematopoietic) and CD43-(endothelial and mesenchymal) cells at different time points during EHT using EdU incorporation assay (**Fig. 4b**). These experiments confirmed that the emerging CD43+ haematopoietic population was characterised by an active cell cycle profile, while the majority of CD43-cells were in the G1 phase. Furthermore, we took advantage of FUCCI-hPSCs (38, 39) to monitor changes in cell cycle phase during EHT (**Supplementary Fig. 6c)**. We observed that emerging HPC clusters contained cells that mainly expressed the reporter specific for S/G2/M phases (GEMININ), while endothelial cells were found to be either positive for the late G1 reporter (CDT1) or negative for both reporters thus indicating G0/early G1 phase (**Supplementary Fig. 6c)**. These findings support the notion that the endothelial differentiation into HPCs is associated with cell cycle entry.

We next asked whether RUNX1+ haemogenic endothelial cells present at EHT D3 could transit to haematopoietic progenitors if their cell cycle progression was blocked. To address this question, EHT D3 cells were grown for 48 hours in the presence of Nocodazole, a small molecule commonly used to block cell cycle progression (40, 41). The efficacy of Nocodazole treatment was confirmed by EdU incorporation analyses showing an expected enrichment of cells in G2/M (**Fig. 4c**). However, a significant fraction of cells remained in G1. These cells most likely represent a quiescent population of endothelial cells (**Fig. 4a**) that never entered cell cycle during the 48 hours treatment. Following Nocodazole treatment, cells were washed to allow cell cycle progression and their haematopoietic potential was assessed by CFU assays. This showed that cell cycle inhibition effectively blocked the generation of functional haematopoietic cells (**Fig. 4d**). Consistently, flow cytometry analyses showed an enrichment in CDH5+ (endothelial) cells and depletion of the CD43+ (haematopoietic) compartment after Nocodazole treatment (**Fig. 4e**). Importantly, this effect was not related to nonspecific cytotoxicity, as cells treated with Nocodazole only displayed a minor increase in cell death (**Supplementary Fig. 6b**).

To confirm that blocking cell cycle does not affect the ability of endothelial cells to subsequently re-enter cell cycle and proliferate, but it rather influences their cell fate decision by precluding the haematopoietic fate choice, we performed release experiments. Following 48 hours Nocodazole treatment from EHT D3 to EHT D5, cells were washed/released and further cultured for up to one week. Following release from cell cycle block, endothelial cells were able to re-enter cell cycle and resume proliferation, expanding as a monolayer, possibly composed of a mix of endothelial and mesenchymal cells. Importantly, immunofluorescence staining confirmed the progressive reduction in the number of RUNX1+ cells (**Fig. 5a**). The rare RUNX1+ cells detectable after treatment did not form distinct clusters, displayed an elongated endothelial morphology, as assessed by immunofluorescence and light microscopy, and were unable to acquire colony forming potential (**Fig. 5b**). In line with these results, there were no visible HPC clusters even after one additional week of culture.

**Figure 5.**
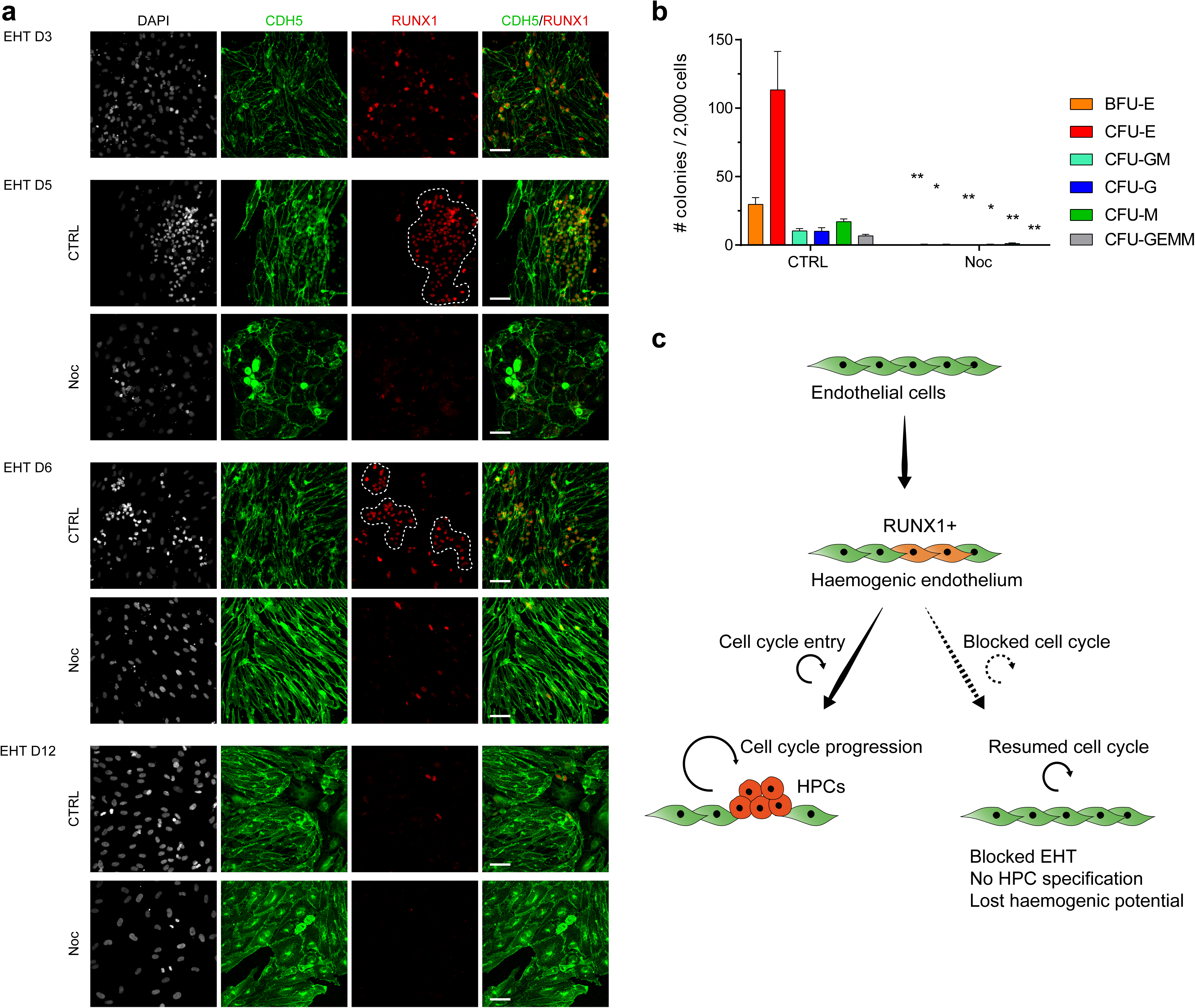
Endothelial cells lose haemogenic potential upon cell cycle block. (A) Immunofluorescence staining showing the expression of CDH5 and RUNX1 over time. At EHT D3, media was supplemented either with 0.1% DMSO (CTRL) or with 0.1 μg/mL Nocodazole (Noc). After 48h, at EHT D5 cells were washed and media refreshed every other day. After cell cycle block is released, cells reprise proliferation, but RUNX1 expression is gradually lost, with positive cells representing very rare events at EHT D12. Haematopoietic clusters are outlined in CTRL and consist of RUNX1+ cells that are mainly round shaped and grouped in clusters. These clusters can also be recognised in the CDH5 stained panel (green), in which they display a typical round morphology, as opposed to the surrounding elongated endothelial cells. Finally, these clusters can also be recognised in the DAPI (grey) panel, in which they are characterised by the typical dense nuclear morphology and brighter appearance compared to endothelial cells. In Nocodazole treated cultures, all these features are not present: the few RUNX1+ cells are more scattered and not grouped in clusters; they mark elongated endothelial cells rather than round cells; these elongated cells are CDH5 positive (green panel) and thus clearly endothelial; finally, their nuclei do not appear to be denser or brighter than the surrounding cells, as was the case in haematopoietic clusters found in control. Scale bar is 50 μm. **(B)** 48h after Nocodazole release, at EHT D7 cells were collected and cultured in a CFU assay to assess their differentiation potential. Data are mean ±SEM of 3 independent experiments, with statistical significance compared to control as *P<0.5, **P<0.01, ***P<0.001, ****P<0.0001 by two-tailed unpaired t-test. Degrees of freedom=4, t-values for the distinct colonies are BFU-E=5.803, CFU-E=4.02, CFU-GM=6.2, CFU-G=3.625, CFU-M=7.407, CFU-GEMM=5.547. **(C)** Diagram summarising the role of cell cycle progression during EHT.

Taken together, these experiments confirmed that blocking cell cycle prevents transition of haemogenic endothelial cells into haematopoietic progenitors (**Fig. 5c**). Furthermore, when cell cycle is blocked, haemogenic endothelial cells progressively lose the expression of RUNX1 and adopt a non-haemogenic endothelial identity or possibly assume a mesenchymal fate.

### Inhibition of specific cyclin-dependent kinase (CDK) proteins impairs EHT

To further explore the connection between cell cycle regulation and haematopoietic differentiation we focused on CDK proteins and their role during EHT. CDK proteins are key regulators of cell cycle progression and are involved in a variety of non-canonical functions such as transcriptional activation, DNA damage repair, metabolism, differentiation and cell fate decision in multiple systems (39, 42, 43, 44, 45). To this end, we used three well-characterised inhibitors which bind to CDK proteins and block their phosphorylation activity (PD0332991 for CDK4 and CDK6; RO3306 for CDK1, and Roscovitine for CDK2 and to a minor extent CDK1), (46, 47, 48, 49, 50, 51, 52, 53).

In short, EHT D3 cells were grown in the presence of each inhibitor for 48 hours and were washed and collected at EHT D5 for further analyses. Specifically, we assessed the cell cycle profile of unenriched populations by EdU staining, and the colony forming capacity of sorted HPCs (**Fig. 6**). Consistent with the known functional redundancy between CDKs, the treatments were not able to completely block cell cycle but instead induced enrichment in specific phases. Thus, this approach was fundamentally different compared to the use of Nocodazole, as it allowed us to examine the role of distinct cell cycle regulators rather than cell cycle progression. More precisely, inhibition of CDK4/6 (CDK4/6i) increased the number of cells in G1, Roscovitine (CDK2/1i) did not significantly change cell cycle profile, inhibition of CDK1 (CDK1i) caused an increase in the number of cells in G2/M (**Fig. 6a**).

**Figure 6.**
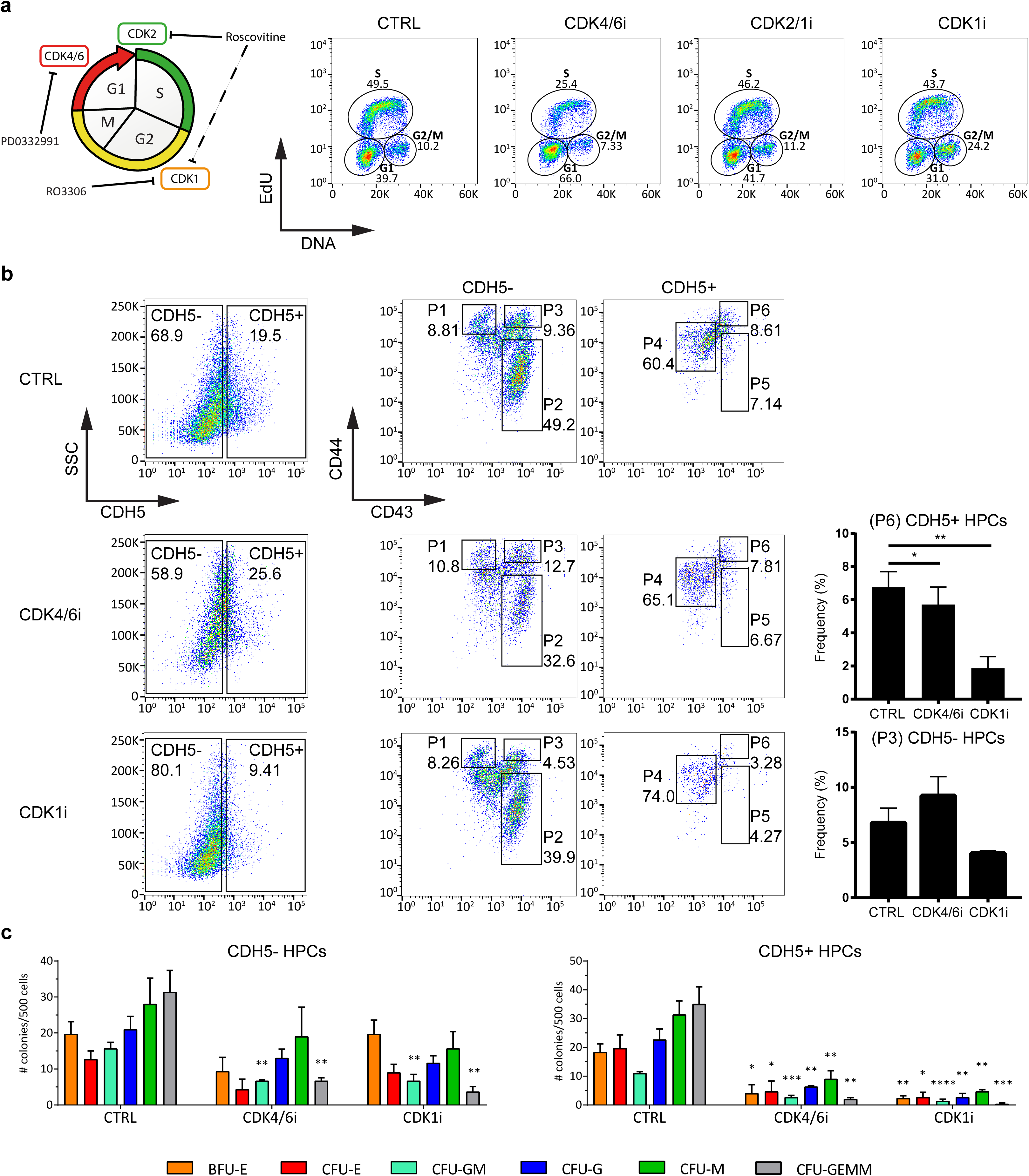
Functional effect of CDK inhibitors during EHT. (A) Schematic showing inhibitors used in the experiments (left panel) and their effect on cell cycle progression (right panel). Samples were treated for 48 hours between EHT D3 and EHT D5 with either 0.1% DMSO (CTRL), 1 μM PD0332991 (CDK4/6i), 10 μM RO3306 (CDK1i) or 4 μM Roscovitine (CDK2/1i). At EHT D5 cells were either **(A)** processed for EdU staining and flow cytometry to monitor cell cycle profile or **(B)** stained for flow cytometry to monitor the frequency of distinct cell populations generated (P1: mesenchymal cells, P4: endothelial cells, P2 and P5: erythroid progenitors, P3 and P6: HPCs). Results are representative of 3 independent experiments. **(C)** At EHT D5 cells were washed to remove the inhibitors and two populations of HPCs (corresponding to P3 and P6 in (B)) were FACS-sorted and further cultured in a CFU assay to reveal their differentiation potential. Data are mean ±SEM of 3 independent experiments, with statistical significance compared to control as *P<0.5, **P<0.01, ***P<0.001, ****P<0.0001 by one-way ANOVA. Degrees of freedom in (B) for CDH5-HPCs are DFn=1.103, DFd=2.205, F value 11.19, for CDH5+ HPCs are DFn=1.046, DFd=2.092, F value 204.2. Degrees of freedom in (C) are DFn=2, DFd=6, F values for the distinct colonies in CDH5-HPCs are BFU-E=2.483, CFU-E=2.771, CFU-GM=12.15, CFU-G=3.261, CFU-M=0.8633, CFU-GEMM=17.43, F values for the distinct colonies in CDH5+ HPCs are BFU-E=12.51, CFU-E=6.757, CFU-GM=67.36, CFU-G=21.71, CFU-M=19.26, CFU-GEMM=30.63.

Further analysis revealed that CDK4/6i treatment resulted in a reduction in the number of CD43+ cells and a modest effect on the frequency of HPCs (**Fig. 6b**). However, the differentiation potential of both CDH5+ and CDH5-HPCs was disrupted since their capacity to form blood colonies was significantly reduced (**Fig. 6c**). Roscovitine was not associated with major modifications in the frequency of populations generated during EHT. Finally, CDK1i significantly decreased the number of HPCs and to a minor extent the number of erythroid CD43+ cells (**Fig. 6b**). The smaller fraction of HPCs produced displayed a strong impairment in their differentiation potential as shown by CFU assays (**Fig. 6c**). Similarly, the inducible knockdown of CDK1 depleted the colony forming potential of the EHT culture, thus confirming our results using a genetic approach (**Supplementary Fig. 7**).

Taken together, these experiments suggest that specific CDKs are necessary not only for EHT to occur but also for HPCs to establish their full differentiation potential. Importantly, the inhibition of distinct cell cycle regulators, as opposed to cell cycle block, alters, rather than inhibit, the EHT process leading to HPC specification.

### Inhibition of CDK regulators disrupts EHT

In order to further understand the effect of CDK inhibitors on the EHT process and the differentiation capacity of HPCs, we performed scRNA-seq analyses on EHT D5 cells grown in the presence of each CDK inhibitor for 48 hours. Following quality control, we analysed 1,742 cells treated with CDK4/6i, 2,140 treated with CDK2/1i and 1,741 with CDK1i. As expected, in each of the samples we identified HPCs, endothelial, mesenchymal and erythroid populations (**Fig. 7a**).

**Figure 7.**
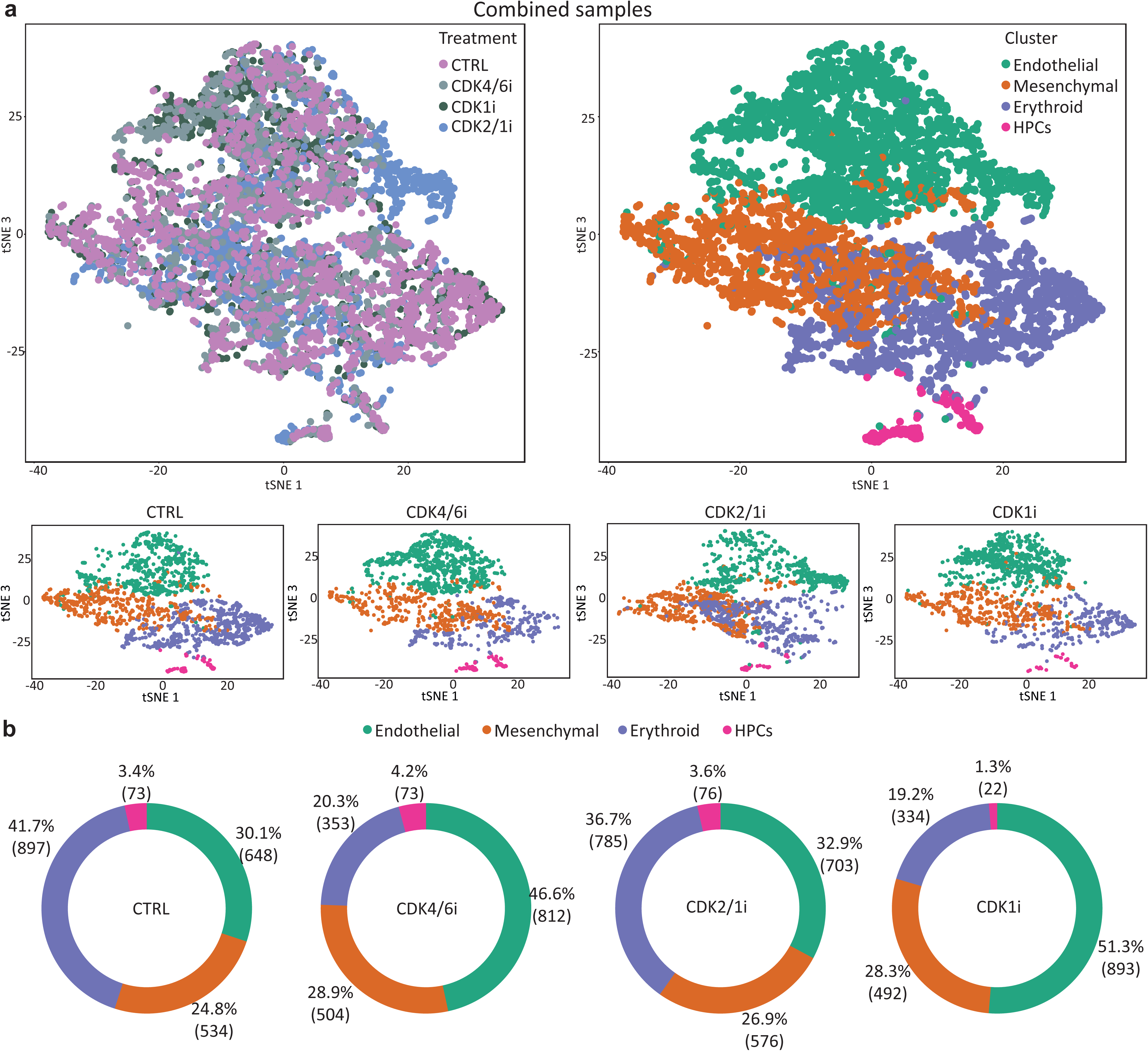
Effect of CDK inhibitors evaluated by single-cell transcriptomics. Samples were treated with different cell cycle inhibitors, namely PD0332991 (CDK4/6i), Roscovitine (CDK2/1i) and R03306 (CDK1i), or with DMSO (CTRL). At EHT D5 cells were collected and processed for scRNA-seq. Data from the four samples was merged and clustered. **(A)** tSNE plots showing previously identified cell populations, on the merged samples (top row) and for each treatment separately (bottom row). **(B)** Pie charts showing the percentage and absolute number of cells for each condition.

Comparative analysis showed increase in the percentage of endothelial cells upon inhibition of CDK4/6, associated with a decrease in the erythroid compartment. Endothelial cells are mostly non-cycling cells (**Fig. 4a, b**, **Supplementary Fig. 6a, c**), and in control conditions their numbers decrease during EHT concomitantly with production of haematopoietic cells (**Fig. 2b**). Thus, our data suggest that the increase in endothelial cells following CDK4/6 inhibition, compared to control, is not associated with proliferation but rather with their decreased transition to the haematopoietic fate, *i.e.* decreased EHT (**Fig. 7b**). The percentage of HPCs in the presence of the inhibitor was comparable to control, confirming our flow cytometry data (**Fig. 6b**) However, these HPCs displayed impaired differentiation potential (**Fig. 6c**). This could explain the reduction in the erythroid compartment, and why the frequency of HPCs remained comparable to control despite reduced EHT.

Cells grown in the presence of the CDK1i also displayed increase in the percentage of endothelial cells and reduction in the erythroid cluster, suggesting a reduced progression through EHT. However, the number of HPCs was strongly reduced when compared to control, again in agreement with our previous flow cytometry analyses (**Fig. 6b**). This indicates a key role for CDK1 activity in the HPC population.

Next, we inferred the cell cycle state by transcriptome analysis of cells in each of the clusters across all treatments. Inhibition of CDK4/6 caused an expected enrichment in G1 of all four populations (**Fig. 8a**). On the other hand, the inhibition of CDK1 was associated with the expected enrichment in G2/M in endothelial and mesenchymal clusters (**Fig. 8a**). Endothelial cells blocked in this phase of the cell cycle most probably represent those that were undergoing either EHT or EndoMT at the start of the treatment. Indeed, while the majority of endothelial cells are in G1, those that are transitioning to haematopoietic and mesenchymal fates are associated with re-entry into the cell cycle and display downregulation of cell cycle inhibitors concomitant with upregulation of proliferation markers (**Fig. 4a, b**, **Supplementary Fig. 3**). Despite the presence of the CDK1 inhibitor, few HPCs were found in S/G2/M, indicating that the treatment either caused HPCs to further differentiate to the erythroid compartment or it affected their survival. Either options (*i.e.*, differentiation to erythroid progenitors or cell death of HPCs upon CDK1 inhibition) could account for the decrease in the HPC cluster compared to pre-treatment levels, and for the fact that the remaining HPCs were mostly in G1 phase, *i.e.* they were not cycling at the time of the treatment and were not affected by the inhibition of CDK1 which gets activated in G2/M. Importantly, the few remaining progenitors, enriched in G1, also displayed a depleted differentiation potential, similarly to HPCs enriched in G1 by the CDK4/6i treatment (**Fig. 6c**). This suggests that the timely activation of these cell cycle regulators is required for the acquisition of haematopoietic differentiation potential.

**Figure 8.**
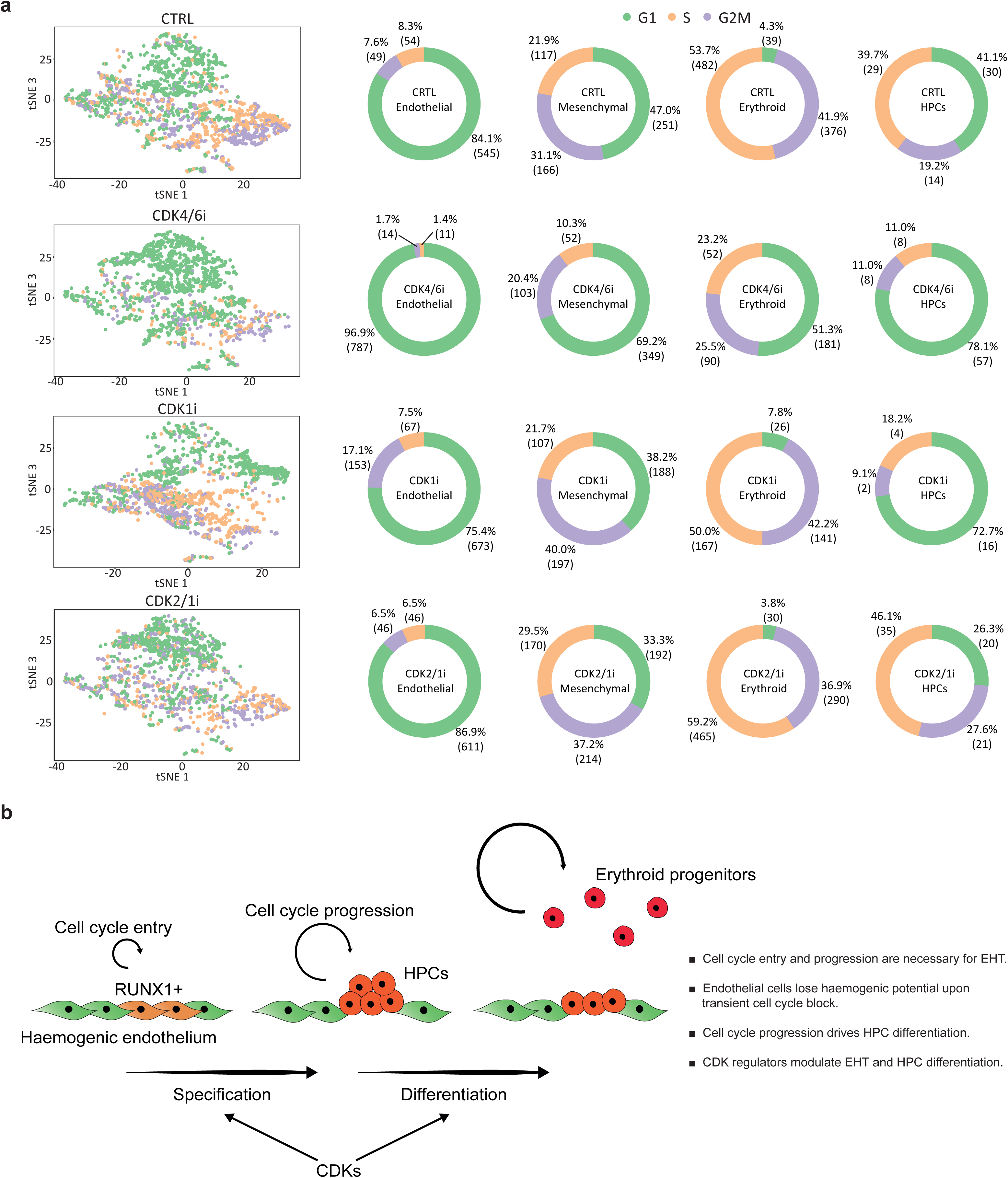
Cell cycle regulation controls EHT. (A) Cell cycle marker genes identified in the scRNA-seq dataset were used to determine the cell cycle profile of individual clusters upon inhibition of distinct CDKs. **(B)** Proposed model for the role of cell cycle during EHT. Endothelial cells need to re-enter the cell cycle in order to progress through EHT and generate functional HPCs. Additionally, endothelial cells lose their haemogenic potential if cell cycle entry and progression is prevented upon transient cell cycle block. The process is regulated by CDKs, which are necessary for EHT and for the acquisition of HPC differentiation potential.

Overall, along with further confirming its requirement for EHT, these results show that cell cycle regulation controls HPC differentiation. Additionally, the two regulators possibly play distinct roles in HPCs. CDK4/6 might be necessary for the acquisition of differentiation potential, as its inhibition preserved the HPC population although negatively affecting its ability to differentiate. CDK1 might be more important for maintaining the HPC population, as the population was eroded when CDK1 was inhibited and only non-cycling cells were able to preserve the HPC state.

Of note, inhibition of either CDK4/6 or CDK1 resulted in modest but broad transcriptional changes across all cell populations compared to control. We mainly observed downregulation of genes for ribosomal subunits and enrichment in GO terms involved in mitochondrial functioning (**Supplementary Fig. 8**, **Supplementary Table 1– 8**). Furthermore, HPCs showed upregulation of genes such as *ALAS2* or haemoglobin genes, suggesting their priming or precocious differentiation toward the erythroid lineage. Importantly, the inhibition of CDK1 was able to affect the transcriptional signature of the HPC population despite not affecting its cell cycle profile, confirming that this regulator may play a role during EHT beyond cell cycle regulation. Unlike CDK4/6 and CDK1, the CDK2/1 inhibitor did not cause major transcriptional changes compared to control, consistent with the fact that this treatment did not affect the EHT process.

To conclude, our single-cell analyses indicate that the inhibition of CDK4/6 and CDK1 were able to cause transcriptional changes that were not strictly related to cell cycle progression. Despite the modest extent of these changes, our findings suggest that these CDKs can participate in a complex crosstalk between cell cycle regulation and the transcriptional programmes that drive EHT.

### *In vitro* EHT culture recapitulates populations found in the human AGM *in vivo*

In order to assess the relevance of our findings for human foetal blood development, we compared the cells generated in our study with a publicly available dataset obtained from human AGM (64). Having annotated this data according to the Authors’ original cell annotations, we then mapped this dataset onto our data. This allowed us to obtain predictions of the cell identities in our data using the annotations from the *in vivo* generated dataset (**Supplementary Fig. 10a**). This approach confirmed that endothelial, mesenchymal and haematopoietic cells identified in our study displayed similarity with corresponding populations described *in vivo*.

Furthermore, endothelial cells generated in our system expressed top marker genes of both haemogenic and arterial endothelium from AGM (**Supplementary Fig. 10b**). Based on the level of expression of marker genes, *in vitro* generated endothelial cells displayed higher similarity to late haemogenic endothelial cells (late HECs), which give rise to true HSPCs in the AGM (64), rather than early haemogenic endothelial cells (early HECs). Interestingly, *in vitro* derived HPCs displayed similarities to end stage HSPCs from AGM, albeit maintaining expression of genes expressed in immature HSPCs such as active cell cycle genes (**Supplementary Fig. 10b**).

Overall, our comparative analysis showed that *in vitro* generated endothelial and haematopoietic cells share many similarities to populations found in the human AGM and thus make a valuable model system for studying EHT.

## DISCUSSION

In this study, we have uncovered a link between cell cycle regulation and endothelial-to-haematopoietic transition (**Figure 8b**). By combining a hPSC differentiation system with single-cell transcriptomics, we gained insights into the mechanisms driving early human haematopoietic development.

Our data suggest that the function of cell cycle regulators goes beyond their canonical role in proliferation (**Supplementary Fig. 9**). Thus, CDK4/6 appear to be necessary for both specification of HPCs (*i.e.* EHT) and their further differentiation. In agreement with our results, genetic studies in the mouse have shown that the cyclin D-CDK4/6 complex is indeed required for haematopoietic development (37, 58). CDK1 also appears to be necessary for EHT, and for balanced HPC differentiation and survival. Of note, CDKs are known to be involved in a variety of non-canonical functions such as transcriptional control and cell fate decision (39, 42, 43, 44, 45, 68). Accordingly, our results showed that the activation of CDK4/6 and CDK1 is required during EHT. Their inhibition was able to affect the transcriptional signature of the distinct populations generated in culture as well as the differentiation capacity of nascent HPCs, further suggesting that the role of these regulators during EHT is not strictly related to cell cycle progression.

Based on these results, we propose a tight link between cell cycle regulation and EHT (**Fig. 8b**). Our model suggests that haemogenic endothelial cells are a population of quiescent or slow proliferating cells. Their transition to the haematopoietic fate is associated with cell cycle re-entry and the acquisition of an active cell cycle profile. This ultimately results in the specification of multipotent HPCs, a transitory population of cells which in the current culture conditions quickly commit to highly proliferative erythroid progenitors. We have shown that cell cycle progression is necessary for EHT, and that upon blocking cell cycle, haemogenic endothelial cells are unable to transition to the haematopoietic fate. As a result, they downregulate RUNX1, lose haemogenic potential and instead adopt a non-haemogenic endothelial cell identity, even when the cell cycle block is removed (**Fig. 5c**, **Supplementary Fig. 9a**). This would suggest that EHT can occur within a narrow time window during which cell cycle entry and progression are necessary for the completion of the haemogenic programme. This is possibly modulated by the timely activation of specific cell cycle regulators.

Recent studies in the mouse have demonstrated that the generation of early haematopoietic progenitors in the AGM is associated with a dynamic cell cycle regulation (59). Our analysis here goes beyond these initial findings and shows that the endothelial cells express key cell cycle inhibitors of the Rb and CIP/KIP families, such as *RB1*, *p21* and *p57* (**Supplementary Fig. 3a**). The downregulation of these inhibitors and concomitant expression of key haematopoietic markers in a subset of endothelial cells mark their re-entry into the cell cycle and the onset of haematopoietic specification. Importantly, blocking cell cycle progression effectively prevents this transition, suggesting that cell cycle could be necessary to orchestrate the expression of key lineage regulators and thus differentiation mechanisms.

Interestingly, our group has previously shown that hPSCs acquire responsiveness to distinct differentiation signals during specific phases of the cell cycle, through a post-translational modulation of SMAD2/3 transcriptional activity controlled by cyclin D-CDK4/6 (**39**). We hypothesise that a similar regulation could occur during EHT and that CDKs could be required for the acquisition of functional haematopoietic identity. Accordingly, it was previously reported that CDK6 can bind to RUNX1 and interfere with its transcriptional activity (60). RUNX1 is an essential regulator of EHT, which synergises with other key factors such as TAL1 and GATA2, and is necessary for the induction of the haematopoietic transcriptional programme. Thus, it is possible that an important role for the cell cycle machinery is to modulate the timely activation of the haematopoietic transcriptional network. Importantly, our results show that when this modulation is impeded by preventing cell cycle progression and disrupting the activation of the cell cycle machinery, endothelial cells permanently lose their haemogenic identity. Our data also imply that EndoMT and EHT represent two alternative cell fate choices undertaken by endothelial cells during foetal haematopoietic development. The mesenchymal population does not appear to be affected by the perturbations in the cell cycle used in this study. However, commitment towards mesenchymal cells appears to take place at an earlier stage, at EHT D0 or earlier, and to be similarly associated with the expression of cell cycle related genes (**Supplementary Fig. 3b**). This would suggest that despite EndoMT and EHT taking place at slightly different time windows, endothelial cells similarly need to enter and progress through cell cycle in order to differentiate towards both haematopoietic and mesenchymal fates. Further investigations would allow to explore this possibility, and to assess whether blocking cell cycle at an earlier time point might be able to prevent EndoMT and maximise the number of cells available for EHT once this block is lifted.

In conclusion, our study has demonstrated a complex interplay between the molecular machineries that control cell cycle progression and EHT. This is achieved through the activation of specific cell cycle regulators, which define the transcriptional landscape and control the differentiation potential of populations generated during EHT. Further understanding the mechanisms involved in the specification of HSPCs will help to design more effective culture conditions for their *ex vivo* expansion and pave the way to the effective *in vitro* generation of HSPCs with long term engraftment potential.

### METHODS Culture of hPSCs

The hIPSC lines A1AT-RR (**62**) and FSPS13B were maintained as previously described (**63**) on plates coated with 10 μg/ml Vitronectin (Stem Cell Technologies) and cultured in E6 media supplemented with 2 ng/ml TGFβ (R&D) and 25 ng/ml FGF2 (Dr. Marko Hyvönen, Department of Biochemistry, University of Cambridge). Cells were maintained at 37 °C and 5% CO2, and passaged every 5-6 days by dissociation with 0.5 mM EDTA (ThermoFisher Scientific). For coating plates, 10 μg/ml Vitronectin in PBS (ThermoFisher Scientific) was applied for at least 1 hour at room temperature.

### Haematopoietic differentiation

Differentiation was performed at 5% O_2_ by adapting a protocol previously described (**23**). Briefly, a serum-free differentiation (SFD) medium was used, consisting of 75% Iscove’s modified Dulbecco’s medium (ThermoFisher Scientific) and 25% Ham’s F12 medium (ThermoFisher Scientific) supplemented with 1% N2 (ThermoFisher Scientific), 0.5% B27 (ThermoFisher Scientific), 0.05% BSA (Sigma-Aldrich), 1 mM ascorbic acid (2-phospho-L-ascorbic acid trisodium salt, Sigma-Aldrich), 4.5×10^-4^ monothioglycerol (Sigma-Aldrich), 2 mM L-glutamine (ThermoFisher Scientific), 150 μg/ml transferrin (Sigma-Aldrich) and 10 ng/ml penicillin/streptomycin (ThermoFisher Scientific). Undifferentiated cells were dissociated using 0.5 mM EDTA and small aggregates were resuspended in SFD medium supplemented with 10 ng/ml BMP4 (R&D) and cultured as embryoid bodies (EBs) on non-treated plates (Starlab). After 24 hours, an equivalent volume of SFD was added directly on top to not perturb the small EBs, supplemented with final concentrations of 10 ng/ml BMP4 and 5 ng/ml FGF2. At day 2, developing EBs were collected and washed. For this, they were centrifuged 3 minutes at 100 g, gently resuspended in SFD supplemented with 10 ng/ml BMP4, 5 ng/ml FGF2 and 3 μM CHIR99021 (Tocris), and seeded back on the plate. After 24 hours, EBs were again collected and washed, this time by centrifuging at 300 g for 3 minutes, and resuspended in StemPro-34 SFM (ThermoFisher Scientific) supplemented with 1 mM ascorbic acid, 4.5×10-4 monothioglycerol, 2 mM L-glutamine, 150 μg/ml transferrin, 10 ng/ml penicillin/streptomycin (from now on referred to as complete SP34) containing 5 ng/ml FGF2, 15 ng/ml VEGF (Peprotech), 10 ng/ml IL-6 (R&D) and 5 ng/ml IL-11 (R&D), and cultured for 48 hours. At day 5, EBs were again collected, washed and resuspended in complete SP34 with 5 ng/ml FGF2, 15 ng/ml VEGF, 10 ng/ml IL-6, 5 ng/ml IL-11, 50 ng/ml SCF (R&D), 5 ng/ml IGF-1 (R&D), 2 U/ml EPO (R&D) and cultured until day 8. At this stage, EBs were collected and prepared for sorting. For this, they were incubated for 20 minutes at 37 °C with collagenase solution, composed of Advanced DMEM/F12 (ThermoFisher Scientific), 20% KSR (ThermoFisher Scientific), 1% L-glutamine and 1 mg/ml collagenase IV (ThermoFisher Scientific), followed by 3 minutes incubation with TrypLE (ThermoFisher Scientific). Single cells were then stained for flow cytometry and the CD34+/CD43-fraction was sorted and used for the second stage of differentiation. For this, the population was resuspended at a density of 10^6^ cells/ml in complete SP34 with 10 ng/ml BMP4, 5 ng/ml FGF2, 5 ng/ml VEGF, 10 ng/ml IL-6, 5 ng/ml IL-11, 100 ng/ml SCF, 25 ng/ml IGF-1, 30 ng/ml TPO (Peprotech), 30 ng/ml IL-3 (Peprotech), 10 ng/ml Flt3-L (R&D), 20 ng/ml SHH (R&D), 10 ng/ml angiotensin II (Sigma-Aldritch) and 100 μM losartan potassium (R&D). Cells were transferred to a non-treated round-bottom 96 well plate (Corning), 200 μl/well (corresponding to 2×10^5^ cells/well), centrifuged 3 minutes at 300 g, and incubated overnight to allow the cells to re-aggregate. On the following day, marking EHT D1, the small aggregates were plated on Matrigel (Corning). For this, they were gently transferred to thin-layer Matrigel-coated wells, with a density of 2×10^5^ cells/well in a 24 well plate, and cultured for additional 2-4 days using the same media, replaced every 2 days. For coating plates, Matrigel was diluted into cold medium with a concentration of 35 μg/cm2 of surface to be coated, and applied overnight at 37 °C. For treatments with cell cycle inhibitors, 0.1 μg/mL Nocodazole (Sigma-Aldrich), 1 μM PD0332991 (Tocris), 4 μM Roscovitine (Sigma-Aldrich), 10 μM RO3306 (Sigma-Aldrich) or 0.1% DMSO were added at EHT D3 for 48 hours.

### Isolation of CD34+ peripheral blood mononuclear cells

Under sterile conditions, peripheral blood was diluted with room temperature PBS supplemented with 1 M trisodium citrate and 20% human serum albumin. 2 volumes of blood dilution were transferred in 50 ml tubes on a layer of 1 volume of Ficoll-Paque (Sigma-Aldrich). After centrifugation for 15 minutes at 800 g, the resulting layer of mononuclear cells was carefully removed, transferred to a new tube, further diluted using the same buffer and centrifuged for 6 minutes at 600 g. The resulting supernatant was removed, the pellet was resuspended in cold PBS supplemented with 0.5 M EDTA and 20% human serum albumin, and centrifuged again for 6 minutes at 600 g. Cells were then processed for CD34+ enrichment using labelling with magnetic beads. Briefly, cells were resuspended in the same PBS/EDTA/human serum albumin buffer, added with 50 μl/108 cells of FcR blocking reagent (Miltenyi Biotec) and 50 μl/10^8^ cells of CD34 magnetic beads (Miltenyi Biotec) and incubated for 30 minutes at 4 °C. After incubation, cells were washed using the same buffer and processed with AutoMACS Pro Separator to enrich for CD34+ peripheral blood mononuclear cells.

### May–Grünwald-Giemsa staining

Cells were resuspended in culture media and concentrated by cytospin centrifugation at 700 g for 5 minutes onto SuperFrostPlus slides (ThermoFisher Scientific) using a Shandon Cytospin 3 cytocentrifuge. Slides were fixed for 3 minutes in cold methanol and stained with May-Grünwald Giemsa (Sigma). Images were captured using a Leica DM5000b microscope in conjunction with a ×63 oil-immersion lens and an Olympus DP72 camera

### Flow cytometry

Cells were dissociated into single cells using TrypLE for 3 minutes at 37 °C, washed with 0.1% BSA-PBS and either fixed with 1% paraformaldehyde and kept at 4 °C for maximum 1 week or immediately stained for flow cytometry. For the staining, after a wash with PBS, cells were blocked with 10% donkey serum (Bio-Rad) for 30 minutes at room temperature. Cells were then stained with the relevant conjugated antibodies diluted in PBS for 1 hour at room temperature, protected from light. Following two washes with PBS, cells were analysed using the Cyan ADP flow cytometer, or sorted on the BD Influx cell sorter. Data was analysed using the FlowJo VX software.

### Antibodies used for flow cytometry

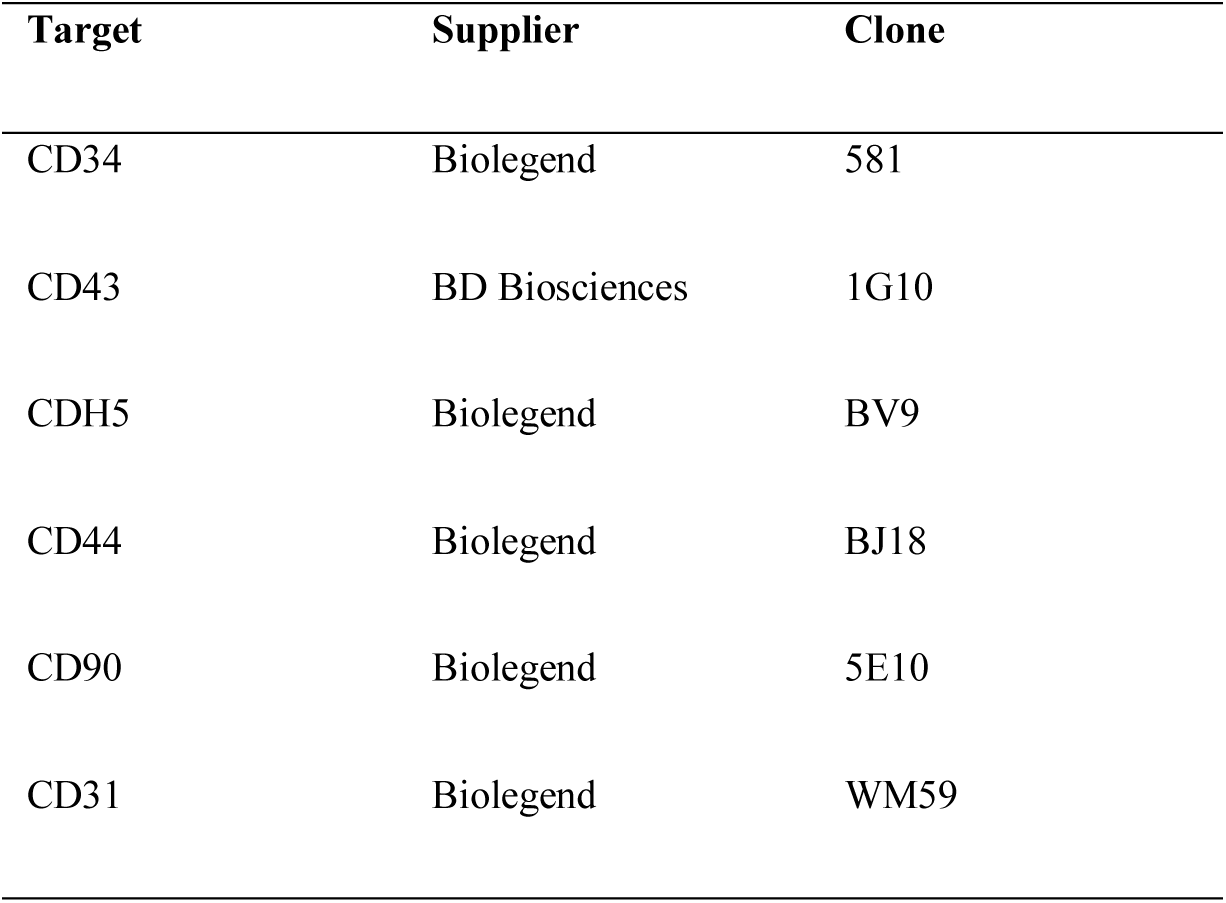

Cell viability upon Nocodazole treatment was assessed using the Dead Cell Apoptosis Kit with Annexin V Alexa Fluor 488 & Propidium Iodide (ThermoFisher Scientific).

### RNA extraction, cDNA synthesis and qPCR

Total RNA was extracted using the GenElute Mammalian Total RNA Miniprep Kit (Sigma-Aldrich) and the On-Column DNase I Digestion set (Sigma-Aldrich) according to the manufacturer’s instructions. RNA was reverse-transcribed using 250 ng random primers (Promega), 0.5 mM dNTPs (Promega), 20 U RNAseOUT (Invitrogen), 0.01 M DTT (Invitrogen) and 25 U of SuperScript II (Invitrogen). For the qPCR reaction, the resulting cDNA was diluted 30-fold. Quantitative PCR mixtures were prepared using the KAPA SYBR FAST qPCR Master Mix Kit (Kapa Biosystems), 4.2 μl of cDNA and 200 nM of each of the forward and reverse primers. Technical duplicates of the samples were run on 384 well plates using the QuantStudio 12K Flex Real-Time PCR System machine and results analysed using the delta cycle threshold (ΔCt) method. Expression values were normalized to the housekeeping gene RPLP0.

### Primers used for qPCR

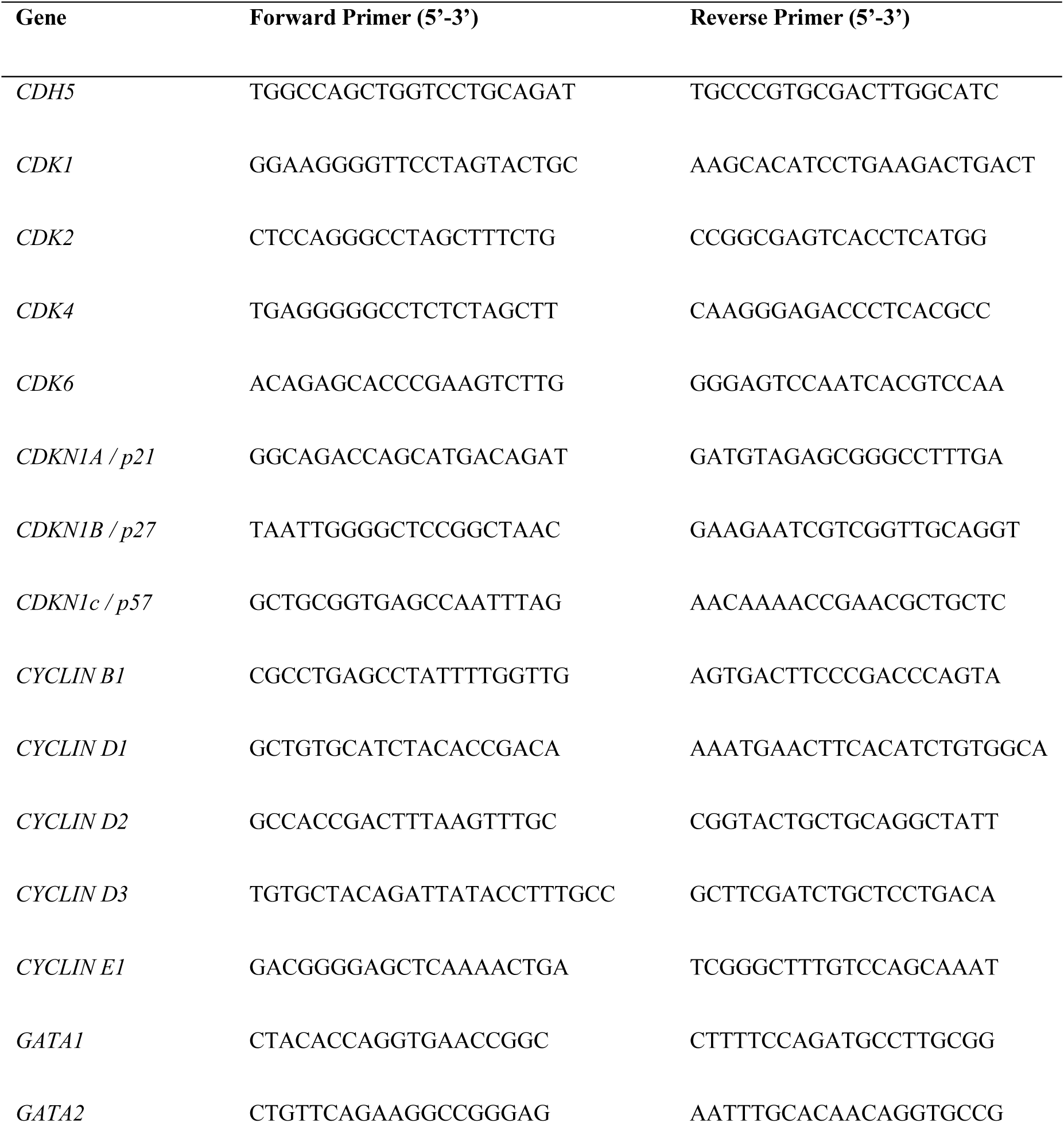
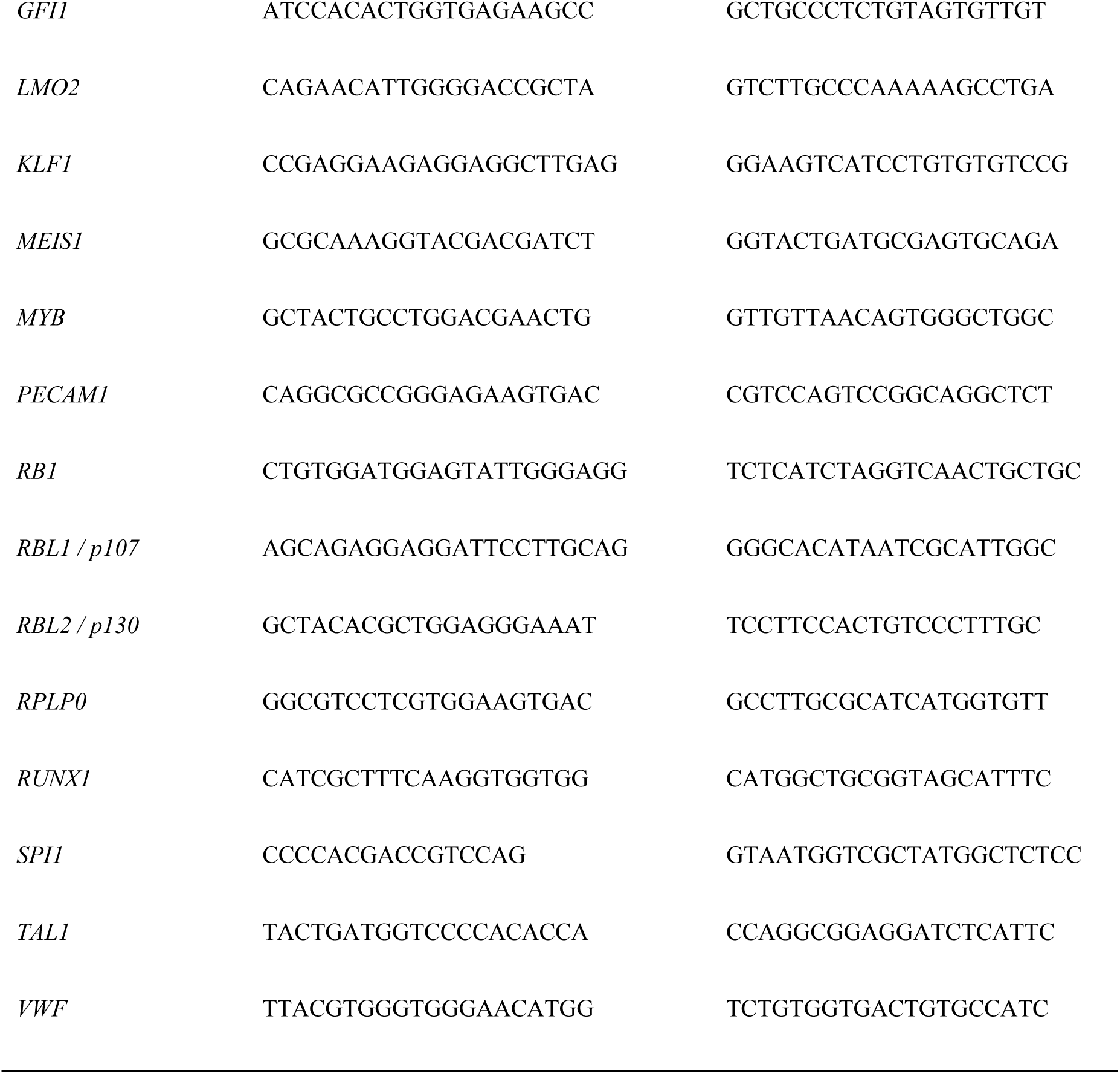

#### CFU assay

The assay was performed using the MethoCult H4435 Enriched medium (STEMCELL Technologies), following the manufacturer’s instructions. Briefly, cells were added in a tube with the medium, mixed by vortex, plated on non-treated 35 mm culture dishes (Corning) and incubated for 14 days at 37 °C, 5% CO2 and 5% O2. Different types of colony were recognised based on morphology, counted and, when relevant, collected for RNA extraction.

#### Cell cycle profile analysis

Cell cycle profile analysis was performed using the Click-iT EdU Alexa Fluor 488 Flow Cytometry Assay Kit (ThermoFisher Scientific) according to the manufacturer’s instructions. Briefly, cultured cells were incubated at 37 °C with 10 μM EdU for 1 hour and harvested after dissociation with TrypLE. After 3 washes with 0.1% BSA-PBS, cells were fixed with 1% paraformaldehyde for 15 minutes at room temperature and washed three more times with 0.1% BSA-PBS. Cells were then permeabilised for 15 minutes with saponin-based permeabilisation/wash buffer and incubated with the Click-iT reaction cocktail for 30 minutes protected from light. Cells were washed once with saponin-based permeabilisation/wash buffer, stained for DNA content using DAPI (ThermoFisher Scientific) and analysed on the Cyan ADP flow cytometer and FlowJo VX software.

#### Immunofluorescence staining

Cells were fixed with 4% paraformaldehyde and kept at 4 °C for maximum 1 week or immediately stained for immunohistochemistry. Cells were blocked/permeabilised with 1% Donkey serum in PBS + 0.3% Triton X-100 for 1 hour, and then incubated with the relevant primary antibodies diluted in PBS + 0.03% Triton X-100 at 4 °C overnight. Cells were then washed three times with PBS and incubated with secondary antibodies diluted in PBS + 0.03% Triton X-100 at 4 °C for 1 hour.

### Antibodies used for immunostaining

| Target | Supplier | Catalogue number |
| --- | --- | --- |
| RUNX1 | Biolegend | 659302 |
| CD43 | BD Biosciences | 560198 |
| CDH5 | R&D | AF938 |
| CD44 | Abcam | Ab157107 |
| VIMENTIN | Dako | M7020 |

#### Inducible knockdown

Plasmid carrying an inducible shRNA was generated as previously described (66). Briefly, shRNA for the CDK1 gene (shCDK1) was obtained from previous publication (67) and introduced by cloning of annealed oligonucleotides in the sOPTiKD plasmid between the BglII and SalI-HF sites.

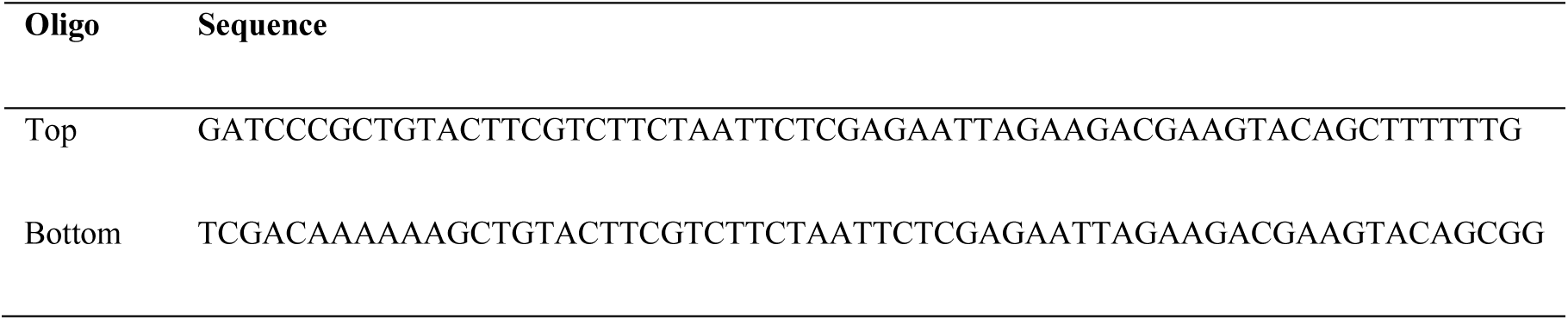

The resulting sOPTiKD_shCDK vector was targeted to the AAVS1 locus in combination with the pZFN.AAVS1-KKR and pZFN.AAVS1-ELD vectors (66). Single cells hPSCs were nucleofected using the Lonza P3 Primary Cell 4D-Nucleofector X Kit and the cycle CA-137 on a Lonza 4D-Nucleofector System, in feeder-free conditions. Monoclonal colonies were selected for 7-10 days with 1 μg/ml Puromycin (Sigma). Expression of the shCDK1 was induced during differentiation by adding 1μg/ml Tetracycline hydrochloride (Sigma-Aldrich) at EHT D2. At EHT D5 the efficiency of knockdown was determined by qPCR, and haematopoietic potential for total culture was assessed by CFU assay.

#### Statistical analysis

Statistical analyses were performed using the GraphPad Prism 7 software. The type of statistical analysis performed and the number of replicates used in each experiment are described in the figure legends. For the comparison of two or multiple groups, two-tailed unpaired t-test or one-way ANOVA test was performed, respectively. Significance in each analysis is represented as *P<0.5, **P<0.01, ***P<0.001, ****P<0.0001.

#### Single cell RNA sequencing

The datasets supporting the conclusions of this article are available in the ArrayExpress repository [E-MTAB-8205]. https://www.ebi.ac.uk/arrayexpress/experiments/E-MTAB-8205/ All the methods were adopted as previously described (26).

#### Single cell RNA processing

Following dissociation, cells were resuspended at a concentration of 1,500 cells/μl in ice-cold SP34 medium, complete with cytokines as for EHT culture. Libraries were constructed using Chromium Controller and Chromium Single Cell 3’ Library & Gel Bead Kit (10x Genomics) according to the manufacturer’s instructions for the recovery of 2,000 cells for each sample.

Briefly, the cellular suspension was added to the master mix containing nuclease-free water, RT Reagent Mix, RT Primer, Additive A and RT Enzyme Mix. Master mix with cells was transferred to the wells in row 1 on the Chromium Single Cell A Chip (10x Genomics). Single Cell 3’ Gel Beads were transferred in row 2 and Partitioning Oil was transferred into row 3.

The chip was loaded on Chromium Controller to generate single cell GEMs. GEM-RT was performed in a C1000 Touch Thermal cycler (Bio-Rad) at the following conditions: 53 °C for 45 minutes, 85 °C for 5 minutes, held at 4 °C. Post GEM-RT clean-up was performed with DynaBeads MyOne Silane Beads (ThermoFisher Scientific). cDNA was amplified using C1000 Touch Thermal cycler at the following conditions: 98 °C for 3 minutes, 12 cycles of (90°C for 15 seconds, 67 °C for 20 seconds and 72 °C for 1 minute), 72 °C for 1 minute, held at 4°C. Amplified cDNA was cleaned with the SPRIselect Reagent Kit (Beckman Coulter) and quality was assessed using 2100 Bioanalyser (Agilent). Libraries were constructed following the manufacturer’s protocol and sequenced in pair-end mode on Hi-Seq4000 platform.

### Alignment and quantification of sequencing data

Cell Ranger v2.10 was used in order to de-multiplex raw base call (BCL) files generated by Illumina sequencers into FASTQ files, perform the alignment, barcode counting and UMI counting. Ensembl BioMart version 91 was used to generate the reference genome.

### Quality control of sequencing data

Data was filtered based on the Median Absolute Deviation (MAD) of the distribution of the number of detected genes. In addition, the percentage of mitochondrial content was set to less than 20%. Following quality control, 1,764 single cells from EHT D0, 3,877 from EHT D3, 2,152 from EHT D5, 1,742 from PD0332991 (CDK4/6i), 2,140 from Roscovitine (CDK2/1i) and 1,741 from RO3306 (CDK1i) were used in downstream analyses.

### Seurat Alignment Strategy

In order to perform a direct comparison of clusters that belonged to the same cell type across different conditions, we adopted the Seurat Alignment workflow (26). We calculated Highly Variable Genes (HVGs) for each of the different conditions. HVGs were detected based on their average expression against their dispersion, by means of the “FindVariableGenes” Seurat command with the following parameters: mean.function equal to ExpMean, dispersion.function equal to LogVMR, x.low.cutoff equal to 0.0125, x.high.cutoff equal to 3, and y.cutoff equal to 0.5. For the analysis of EHT D3 and EHT D5 we selected 1,289 common HVGsthat were expressed in both datasets. For the analysis of EHT D5, CDK4/6i, CDK2/1i and CDK1i we selected 2,306 HVGs that were expressed in at least 2 out of 4 samples. Canonical Correlation Analysis (CCA) was then performed in order to identify shared correlation structures across the different conditions using the “RunMultiCCA” command. 30 and 35 CCA components were selected for the EHT D3 - EHT D5 and the EHT D5 - CDK4/6i - CDK2/1i - CDK1i analysis, respectively, by means of the shared correlation strength, using the “MetageneBicorPlot” Seurat command. Aligned CCA space was then generated with the “AlignSubspace” Seurat command. Significant CCA aligned components were then used to create the 3D tSNE space using the “RunTSNE” Seurat command.

### Downstream analysis of sequencing data

For the clustering in the 3D tSNE space we used the “FindClusters” command in Seurat that performs the Shared Nearest Neighbor (SNN) modularity optimization based clustering algorithm in Correlation Component Analysis (CCA) aligned space. In total, thirteen clusters were identified using SNN modularity optimisation based clustering algorithm on the significant CCA aligned components at 0.6 resolution. Positive marker genes that were expressed in at least half of the cells within the thirteen identified clusters were calculated with “FindAllMarkers” Seurat command, using Wilcoxon rank sum test with the threshold set to 0.25. We over-clustered the cells and then calculated for each cluster the average expression level of the top 20 marker genes. After calculating the correlation across the clusters we merged those with a correlation higher than 0.9. By merging the most highly correlated, we ended up with 4 clusters, and by calculating marker genes we were able to assign cell identity to the resulting clusters.

### Pseudotime ordering

The set of common HVGs (1,289 genes) was used to order cells along a pseudotime trajectory using the Monocle2 R package v1.99.0. Monocle object was created using the ‘importCDS’ command applied to the previous step generated Seurat Object. The pseudotime trajectory was generated by means of the “reduceDimension” command using as additional parameters the‘DDRTree’ reduction method, number of resulted components equal to three, pseudo expression equal to one, using the variance-stabilised matrix of the expression values (norm_method = “log”). Finally, we identified genes that change as a function of pseudotime across each of the three branches by setting the ‘fullModelFormulaStr’ parameter equal to ‘∼sm.ns(Pseudotime)’. For the subclustering, we retrieved only the endothelial cluster Seurat based defined cells (cluster 1) and performed separate louvain clustering on the Monocle space ending up with four subclusters.

### Cell cycle analysis

In order to infer cell cycle states in our single cell data we adopted the Satija’s single cell scoring strategy (26). In more details, we assigned to each cell a score based on its expression of G2/M and S phase markers. These marker sets should be anticorrelated in their expression levels, and cells not expressing either are likely not cycling or in G1 phase. We assigned scores in the ‘CellCycleScoring’ function, which stores S and G2/M scores in ‘object@meta.data’, along with the predicted classification of each cell in either G2M, S or G1 phase.

### Comparative analysis

The GEO dataset GSE135202 derived from human foetal AGM (64) was located and the relevant raw count data from the corresponding single-cell samples GSM3993420:CS10_body_10x, GSM3993421:CS11_CH_10x and GSM3993422:CS13_DA_10x were collected, including their annotation found in Table S3 (Cell annotation and coordinates of early and late populations), related to Figs. 1-4 of the main manuscript. In order to verify that the data were collected properly and the cell type correctly annotated and matched to the publication, we processed the raw counts using FindIntegrationAnchors and IntegrateData Seurat (v3.1) commands, as previously described (65). Having confirmed that cells of similar cell type were clustered together forming well defined clusters in UMAP space, we used the specific dataset as a reference and mapped the defined cell types into our data. This was performed by means of the Referenced Based Integration and Label Transfer Seurat approach (65), using FindTransferAnchors and TransferData Seurat (v3.1) commands. We set as unknown cell type cells that scored less than 30% prediction probability score. Cell types that were present in GSE135202 but not predicted in our tSNE space were also omitted from the plot. This allowed us to represent our control samples in tSNE space coloured based on GSE135202 predictions. Additionally, custom clustering analysis was performed using the Louvain clustering method (FindNeighbors and FindClusters Seurat commands) at 0.3 resolution. Finally, expression of marker genes defining key populations in Figs. 1-4 of the main manuscript were assessed against the cell clusters defined in our study.

### Authors contribution

Conceptualization, G.C., E.A., R.G., L.V., D.O. and A.C.; Methodology, G.C., E.A., R.G., L.V., D.O. and A.C.; Software, E.A.; Formal Analysis, G.C. and E.A.; Investigation, G.C., J.G. and P.S.; Writing –Original Draft, G.C.; Writing –Review & Editing, G.C., L.V., D.O., A.C.; Visualization, G.C., P.S.; Supervision, L.V., D.O., A.C.; Funding acquisition, L.V., A.C.

## Acknowledgment.

This research was supported by a British Heart Foundation PhD Studentship as part of the BHF Oxbridge Centre of Regenerative Medicine (G.C.); Cancer Research UK grant number C45041/A14953 (A.C. and E.A.); European Research Council project 677501 – ZF_Blood (A.C.); the INTENS EU fp8 consortium (D.O); the ERC advanced grant New-Chol (R.G. and L.V.); the Cambridge University Hospitals National Institute for Health Research Biomedical Research Centre (R.G., D.O. and L.V.); and a core support grant from the Wellcome Trust and MRC to the Wellcome Trust – Medical Research Council Cambridge Stem Cell Institute. The work was also supported by the Cambridge NIHR BRC Cell Phenotyping Hub.

## SUPPLEMENTARY FIGURES LEGENDS

**Supplementary figure 1 Characterisation of the differentiation system. (A)** Colonies generated by CFU assay from cells collected at EHT D5. Scale bar as indicated. BFU-E: burst forming unit-erythroid, CFU-E: colony forming unit-erythroid, CFU-GM: colony forming unit-granulocyte-macrophage, CFU-G: colony forming unit-granulocyte, CFU-M: colony forming unit-macrophage, CFU-GEMM: colony forming unit-granulocyte-erythrocyte-macrophage-megakaryocyte. **(B)** qPCR analysis of cells collected after CFU assay, showing the expression of different haemoglobin isoform genes. Data are means ±SEM of 3 independent experiments. **(C)** qPCR analysis of cells collected at different time points during EHT, compared to primary CD34+ cells isolated from peripheral blood. PBMC: peripheral blood mononuclear cells. Data are means ±SEM of 3 independent experiments for EHT samples and 2 independent samples for PBMCs.

**Supplementary figure 2 Expression of lineage marker genes in the scRNA-seq dataset. (A)** Violin plots showing the expression of key lineage markers per cluster. The analysis refers to merged data from EHT D3 and EHT D5 samples. Plots are log2 scaled of the counts. **(B)** May-Grünwald-Giemsa staining of cells collected at EHT D5 and prepared on microscopy slides by cytospin centrifugation. The assay revealed immature cell populations with the absence of obvious morphologies indicating terminally differentiated blood cells in culture, consistent with previous studies of mouse haemogenic endothelium during EHT from E11.5 AGM (61). Scale bar is 15 μm.

**Supplementary figure 3 Dissection of endothelial cell commitment. (A)** Subclustering of the endothelial population. (Left panel) Merged data from EHT D3 and EHT D5 showing differentiation trajectory for endothelial cells only. Cells were coloured to indicate distinct subclusters. (Right panel) Violin plots showing the expression of key lineage and cell cycle genes on endothelial subclusters. Plots are log2 scaled of the counts. All the cells composing the trajectory in this figure are endothelial cells from the endothelial cell cluster (Fig. 2), and violin plots show the expression of different genes in different subsets of the endothelial cluster. **(B)** scRNA-seq analysis of cells sorted at EHT D0. (Left panel) tSNE plot showing cells sorted as CD34+/CD43- and used for the subsequent EHT culture. Six populations were identified based on the transcriptional profiles. (Right panel) Heatmap showing the expression of lineage-associated and cell cycle genes on each cluster. The majority of the cells analysed express genes consistent with an endothelial identity; haematopoietic genes are not expressed at this stage; a small fraction of cells (cluster 4, and to a minor extent cluster 3) co-express endothelial and mesenchymal genes. This also corresponds to the expression of cell cycle related genes.

**Supplementary figure 4 Characterisation and validation of distinct populations. (A)** Immunofluorescence images of samples at EHT D5, showing co-expression of RUNX1 and CD43 (upper panel) or RUNX1 and CDH5 (lower panel) in haematopoietic clusters. Cells showing dim RUNX1 expression but CD43 negative (upper panel) or CDH5 positive (lower panel) possibly represent haemogenic endothelial cells prior to the haematopoietic transition. Scale bar is 50 μm. **(B)** Representative image of endothelial cells (ECs) sorted at EHT D3 as CDH5+/CD44+/CD43-(P4 in Fig. 3) and grown for 10 days. Scale bar is 200 μm. (C) The majority of these cells are positive for the endothelial markers CD31 and CD90, with some of the cells positive for CD34. **(D)** Importantly, these re-plated culture also contains HPCs (P3 and P6), erythroid cells (P2 and P5) and mesenchymal cells (P1), confirming that sorted endothelial cells are able to generate the described populations. **(E)** Representative image of mesenchymal cells (MCs) sorted at EHT D3 as CDH5-/CD44high/CD43-(P1 in Fig. 3) and grown for 7 days. Scale bar is 200 μm. (F) Most of these cells appear negative or low positive for endothelial markers CD31 and CD90, with very few double positive events. **(G)** Immunofluorescence staining demonstrates that these cells are also negative for CDH5, but positive for CD44 and Vimentin (VIM), confirming their mesenchymal identity. Results in B-G are representative of 2 independent experiments.

**Supplementary figure 5 Time course during EHT. (A)** Cells sorted at EHT D0 as CD34+/CD43-(left) were also shown to be CD44low/+ (right). Black box shows corresponding populations in the two panels. **(B)** The sorted population was followed through EHT to monitor the expression of CDH5, CD43 and CD44. This showed that a starting population of CDH5+/CD44low/+ cells gradually acquires brighter expression of CD44 and generate on one side CD43+ cells (HPCs and erythroid), on the other CD44high/CDH5-cells (mesenchymal). Results are representative of 2 independent experiments.

**Supplementary figure 6 EHT is associated with cell cycle entry. (A)** Violin plots showing the expression of key cell cycle regulators in distinct clusters (as defined in Fig. 2). Merged data from EHT D3 and EHT D5, plots are log2 scaled of the counts. **(B)** Apoptosis and cell death staining of Nocodazole treated cultures. At EHT D3, media was supplemented either with 0.1% DMSO (CTRL) or with 0.1 μg/mL Nocodazole (Noc). After 48h, at EHT D5 cells were washed to remove Nocodazole as in previous experiments. Cells were collected after further 48h in culture to detect any long-term toxic effect of the treatments and stained for Annexin V and Propidium Iodide (PI) to monitor apoptotic (Annexin V+) and dead cells (PI+). **(C)** Immunofluorescence images of differentiated FUCCI-hPSCs at EHT D5. The Geminin reporter is expressed during S/G2/M phase and is only expressed in haematopoietic clusters (CD43 positive in upper panel, CDH5 positive and round shaped in lower panel). The CDT1 reporter is expressed during late G1 and is mainly expressed in cells surrounding haematopoietic clusters. Cells in G0/early G1 are negative for both reporters. Scale bar is 50 μm.

**Supplementary figure 7 Validation of the role of CDK1 by inducible knockdown. (A)** Genetic tool that allows the inducible expression of a shRNA upon addition of tetracyclin (TET) in culture for the inducible knockdown of CDK1. **(B)** Cells were treated with TET for 3 days starting at EHT D2. qPCR analysis shows that the induction leads to a 70% reduction of CDK1 mRNA. (C) Cells were either supplemented at EHT D3 with 10 μM RO3306 (CDK1i) for 48h, or at EHT D2 with 0.1% DMSO (CTRL) or 1 μg/ml tetracycline for 72h. At EHT D5, cells were collected, washed and cultured in a CFU assay to assess their differentiation potential. Results are representative of 2 independent experiments.

**Supplementary figure 8 Effect of CDK inhibitors on gene expression in each cluster.** Regression plots showing differentially expressed genes per cluster between treatment and control. For each gene and on each sample, axes are log2 of the average counts. Differentially expressed genes are highlighted. Cut-off used is log2 fold change = 0.5.

**Supplementary figure 9 Proposed model for the role of cell cycle during EHT. (A)** Cell cycle progression is needed for endothelial cells to progress to the haematopoietic fate. If cell cycle is shortly blocked and subsequently released, the haemogenic potential is lost and endothelial cells start proliferating as a monolayer. **(B)** CDK4/6 is important for EHT and for HPC ability to further differentiate. **(C)** CDK1 is important for EHT and for HPC maintenance and survival.

**Supplementary figure 10 scRNA-seq reveals similarities between *in vitro* EHT culture and *in vivo* AGM. (A)** Dataset from human AGM (**64**) was annotated according to Authors’original annotations. These were then used to predict cell identities in our scRNA-seq dataset to detect similarities between *in vitro* and *in vivo* generated cell types. The analysis confirmed that endothelial, mesenchymal and haematopoietic cells identified in our dataset displayed similarity with corresponding populations described in vivo **(B)** Heatmap of the marker genes defining different populations *in vivo*, showing their expression in the clusters originally identified in our dataset. Early/Late HEC: early/late haemogenic endothelial cells, HC: haematopoietic cells, HSPC1-3: haematopoietic stem/progenitor cells type 1-3. Similarly to AGM-derived haemogenic endothelial cells, endothelial cells generated *in vitro* expressed both haemogenic and arterial genes. Additionally, *in vitro* derived endothelial cells displayed higher similarity with late HECs, from which true HSPCs are generated in the AGM, rather than to early HECs. HPCs generated *in vitro* displayed similarities with immature HSPC2 and end stage HSPC3 populations.

## SUPPLEMENTARY TABLES

**Table 1.**
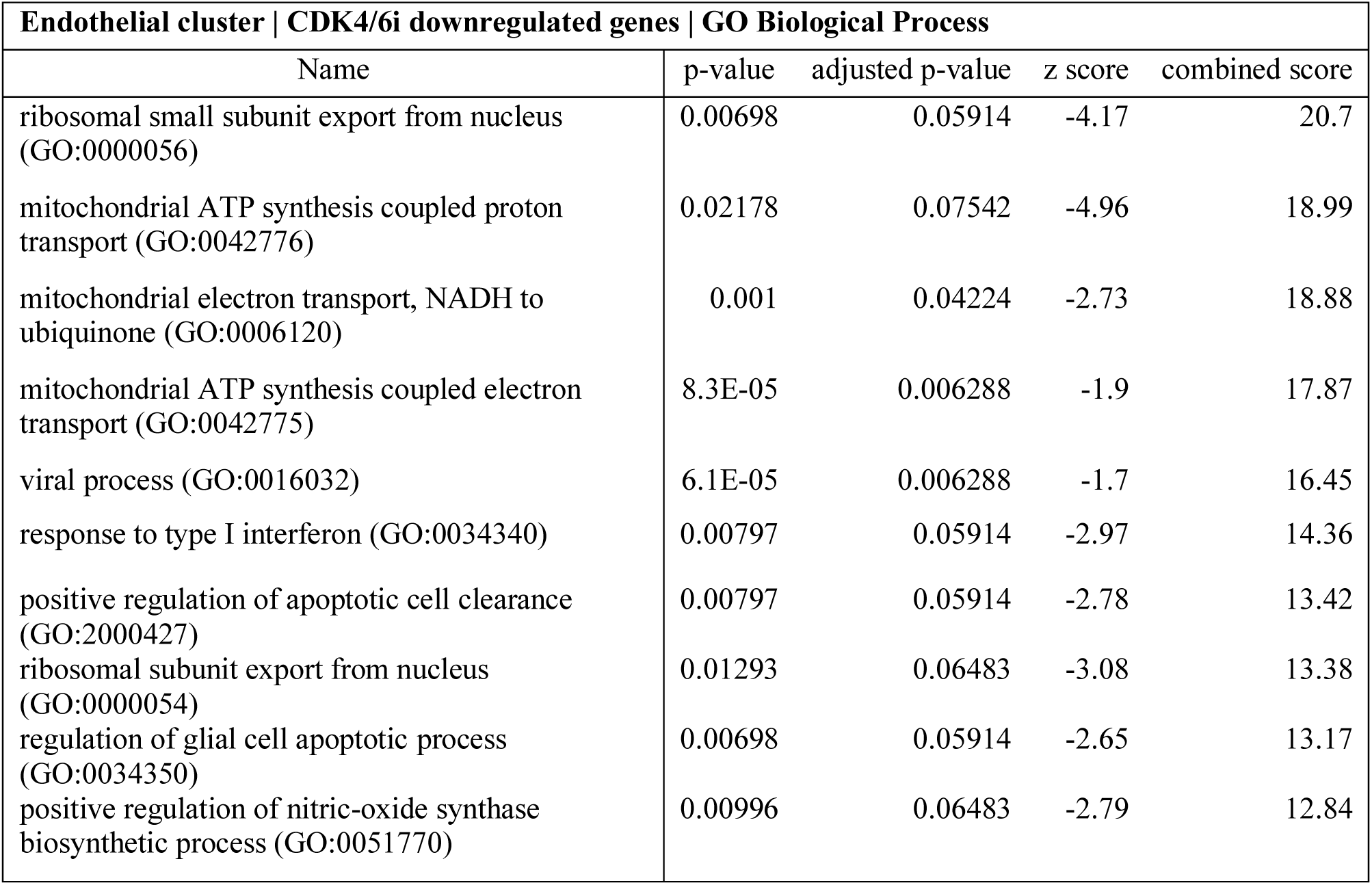
GO terms for CDK4/6i differentially expressed genes in the endothelial cluster.

**Table 2.**
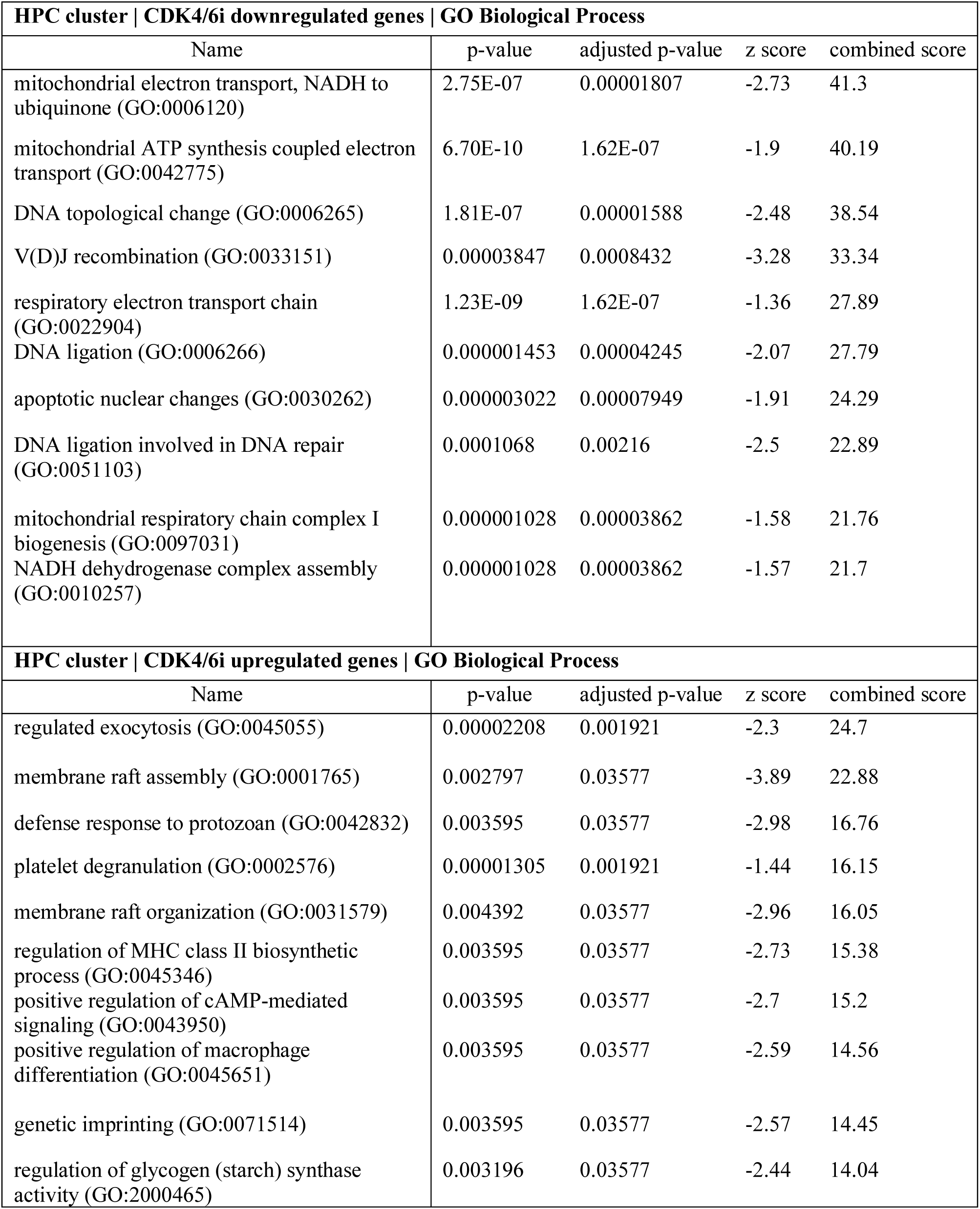
GO terms for CDK4/6i differentially expressed genes in the HPC cluster.

**Table 1.** GO terms for CDK4/6i differentially expressed genes in the erythroid cluster.

| **Erythroid cluster \| CDK4/6i downregulated genes \| GO Biological Process** |  |  |  |  |
| --- | --- | --- | --- | --- |
| Name | p-value | adjusted p-value | z score | combined score |
| mitochondrial ATP synthesis coupled electron transport (GO:0042775) | 2.92E-10 | 1.25E-07 | -1.9 | 41.77 |
| mitochondrial electron transport, NADH to ubiquinone (GO:0006120) | 0.000002073 | 0.0002228 | -2.73 | 35.77 |
| V(D)J recombination (GO:0033151) | 0.000103 | 0.00492 | -3.28 | 30.13 |
| respiratory electron transport chain (GO:0022904) | 5.92E-10 | 1.27E-07 | -1.36 | 28.88 |
| mitochondrial respiratory chain complex assembly (GO:0033108) | 0.000001286 | 0.0001843 | -1.78 | 24.16 |
| DNA topological change (GO:0006265) | 0.0001651 | 0.007099 | -2.47 | 21.54 |
| DNA geometric change (GO:0032392) | 0.00001531 | 0.0008227 | -1.87 | 20.72 |
| DNA ligation involved in DNA repair (GO:0051103) | 0.0002851 | 0.009431 | -2.5 | 20.44 |
| mitochondrial respiratory chain complex I biogenesis (GO:0097031) | 0.000007671 | 0.0004712 | -1.58 | 18.59 |
| NADH dehydrogenase complex assembly (GO:0010257) | 0.000007671 | 0.0004712 | -1.57 | 18.54 |
| **Erythroid cluster \| CDK4/6i upregulated genes \| GO Biological Process** |  |  |  |  |
| Name | p-value | adjusted p-value | z score | combined score |
| membrane raft assembly (GO:0001765) | 0.004193 | 0.05535 | -3.89 | 21.31 |
| regulation of viral entry into host cell (GO:0046596) | 0.0001149 | 0.02343 | -1.89 | 17.14 |
| regulation of muscle system process (GO:0090257) | 0.005985 | 0.05535 | -2.94 | 15.05 |
| membrane raft organization (GO:0031579) | 0.006582 | 0.05535 | -2.96 | 14.86 |
| genetic imprinting (GO:0071514) | 0.005388 | 0.05535 | -2.57 | 13.41 |
| response to muscle stretch (GO:0035994) | 0.005985 | 0.05535 | -2.57 | 13.14 |
| regulation of glycogen (starch) synthase activity (GO:2000465) | 0.004791 | 0.05535 | -2.44 | 13.05 |
| negative regulation of DNA metabolic process (GO:0051053) | 0.0003079 | 0.0314 | -1.52 | 12.28 |
| regulation of apoptotic process (GO:0042981) | 0.001047 | 0.05535 | -1.79 | 12.27 |
| positive regulation of G1/S transition of mitotic cell cycle (GO:1900087) | 0.01016 | 0.05535 | -2.66 | 12.22 |

**Table 4.** GO terms for CDK4/6i differentially expressed genes in the mesenchymal cluster.

| Mesenchymal cluster \| CDK4/6i downregulated genes \| GO Biological Process |  |  |  |  |
| --- | --- | --- | --- | --- |
| Name | p-value | adjusted p-value | z score | combined score |
| mitochondrial electron transport, NADH to ubiquinone (GO:0006120) | 0.000009682 | 0.000568 | -2.74 | 31.58 |
| mitochondrial ATP synthesis coupled electron transport (GO:0042775) | 9.31E-07 | 0.0001222 | -1.9 | 26.42 |
| ribosomal small subunit export from nucleus (GO:0000056) | 0.006284 | 0.04096 | -4.16 | 21.09 |
| mitochondrial ATP synthesis coupled proton transport (GO:0042776) | 0.01962 | 0.05571 | -4.94 | 19.43 |
| respiratory electron transport chain (GO:0022904) | 0.000001389 | 0.0001222 | -1.36 | 18.33 |
| mitochondrial respiratory chain complex I biogenesis (GO:0097031) | 0.00002582 | 0.0007574 | -1.58 | 16.68 |
| NADH dehydrogenase complex assembly (GO:0010257) | 0.00002582 | 0.0007574 | -1.57 | 16.64 |
| mitochondrial respiratory chain complex assembly (GO:0033108) | 0.00008825 | 0.002219 | -1.78 | 16.58 |
| V(D)J recombination (GO:0033151) | 0.007179 | 0.04211 | -3.26 | 16.12 |
| mitochondrial respiratory chain complex I assembly (GO:0032981) | 0.00002582 | 0.0007574 | -1.41 | 14.9 |

**Table 5.** GO terms for CDKli differentially expressed genes in the endothelial cluster.

| **Endothelial cluster \| CDK1i downregulated genes \| GO Biological Process** |  |  |  |  |
| --- | --- | --- | --- | --- |
| Name | p-value | adjusted p-value | z score | combined score |
| positive regulation of nitric-oxide synthase biosynthetic process (GO:0051770) | 0.0000383 | 0.01255 | -2.82 | 28.65 |
| regulation of nitric-oxide synthase biosynthetic process (GO:0051769) | 0.00006628 | 0.01255 | -2.58 | 24.83 |
| ribosomal small subunit export from nucleus (GO:0000056) | 0.006632 | 0.05774 | -4.15 | 20.81 |
| mitochondrial ATP synthesis coupled proton transport (GO:0042776) | 0.0207 | 0.05998 | -4.86 | 18.84 |
| positive regulation of activated T cell proliferation (GO:0042104) | 0.0001608 | 0.01267 | -2.15 | 18.76 |
| positive regulation of cell migration by vascular endothelial growth factor signaling pathway (GO:0038089) | 0.007576 | 0.05774 | -3.47 | 16.95 |
| regulation of positive chemotaxis (GO:0050926) | 0.008519 | 0.05774 | -3.55 | 16.93 |
| protein kinase B signaling (GO:0043491) | 0.0005284 | 0.02313 | -1.97 | 14.87 |
| response to type I interferon (GO:0034340) | 0.007576 | 0.05774 | -2.95 | 14.4 |
| positive regulation of apoptotic cell clearance (GO:2000427) | 0.007576 | 0.05774 | -2.76 | 13.47 |
| **Endothelial cluster \| CDK1i upregulated genes \| GO Biological Process** |  |  |  |  |
| Name | p-value | adjusted p-value | z score | combined score |
| cytoskeleton-dependent intracellular transport (GO:0030705) | 0.00007845 | 0.006058 | -2.09 | 19.73 |
| regulation of transcription from RNA polymerase II promoter in response to oxidative stress (GO:0043619) | 0.004492 | 0.04395 | -3.6 | 19.45 |
| cell cycle G2/M phase transition (GO:0044839) | 0.00002973 | 0.005055 | -1.82 | 18.98 |
| V(D)J recombination (GO:0033151) | 0.003994 | 0.04395 | -3.27 | 18.05 |
| regulation of mitotic cell cycle phase transition (GO:1901990) | 0.00008908 | 0.006058 | -1.91 | 17.85 |
| outer mitochondrial membrane organization (GO:0007008) | 0.003994 | 0.04395 | -3.23 | 17.84 |
| regulation of G2/M transition of mitotic cell cycle (GO:0010389) | 0.00004774 | 0.005411 | -1.77 | 17.61 |
| negative regulation of translation in response to stress (GO:0032055) | 0.003495 | 0.04395 | -3.09 | 17.48 |
| regulation of translational initiation in response to stress (GO:0043558) | 0.003495 | 0.04395 | -2.84 | 16.07 |
| regulation of nuclease activity (GO:0032069) | 0.003495 | 0.04395 | -2.83 | 15.99 |

**Table 6.** GO terms for CDKli differentially expressed genes in the HPC cluster.

| **HPC cluster \| CDK1i downregulated genes \| GO Biological Process** |  |  |  |  |
| --- | --- | --- | --- | --- |
| Name | p-value | adjusted p-value | z score | combined score |
| mitochondrial ATP synthesis coupled electron transport (GO:0042775) | 8.30E-11 | 4.67E-08 | -1.9 | 44.16 |
| mitochondrial electron transport, NADH to ubiquinone (GO:0006120) | 0.000001458 | 0.0001368 | -2.73 | 36.68 |
| respiratory electron transport chain (GO:0022904) | 2.06E-10 | 5.79E-08 | -1.36 | 30.32 |
| mitochondrial ATP synthesis coupled proton transport (GO:0042776) | 0.003724 | 0.03913 | -4.95 | 27.71 |
| mitochondrial electron transport, cytochrome c to oxygen (GO:0006123) | 0.0001003 | 0.002822 | -2.57 | 23.62 |
| mRNA splicing, via spliceosome (GO:0000398) | 0.000001398 | 0.0001368 | -1.75 | 23.54 |
| histone mRNA metabolic process (GO:0008334) | 0.0001484 | 0.003632 | -2.29 | 20.16 |
| gene expression (GO:0010467) | 1.29E-07 | 0.00002419 | -1.26 | 19.94 |
| establishment of protein localization to mitochondrion (GO:0072655) | 0.00005491 | 0.001932 | -2.02 | 19.86 |
| nucleocytoplasmic transport (GO:0006913) | 0.00004293 | 0.001859 | -1.95 | 19.59 |
| **HPC cluster \| CDK1i upregulated genes \| GO Biological Process** |  |  |  |  |
| Name | p-value | adjusted p-value | z score | combined score |
| cellular iron ion homeostasis (GO:0006879) | 0.000003262 | 0.0008498 | -2.65 | 33.52 |
| iron ion homeostasis (GO:0055072) | 9.99E-08 | 0.00005206 | -1.57 | 25.36 |
| oxygen homeostasis (GO:0032364) | 0.01253 | 0.09524 | -4.65 | 20.36 |
| positive regulation of cell cycle arrest (GO:0071158) | 0.00001468 | 0.001912 | -1.71 | 19.01 |
| ribosomal small subunit export from nucleus (GO:0000056) | 0.01253 | 0.09524 | -4.12 | 18.06 |
| gas homeostasis (GO:0033483) | 0.01431 | 0.09524 | -4.11 | 17.47 |
| neutrophil degranulation (GO:0043312) | 0.0001957 | 0.01014 | -2.01 | 17.14 |
| cellular transition metal ion homeostasis (GO:0046916) | 0.00002402 | 0.002503 | -1.59 | 16.88 |
| neutrophil mediated immunity (GO:0002446) | 0.000214 | 0.01014 | -1.93 | 16.31 |
| DNA damage response, signal transduction by p53 class mediator (GO:0030330) | 0.00001468 | 0.001912 | -1.42 | 15.84 |

**Table 7.** GO terms for CDKli differentially expressed genes in the erythroid cluster.

| **Erythroid cluster \| CDK1i downregulated genes \| GO Biological Process** |  |  |  |  |
| --- | --- | --- | --- | --- |
| Name | p-value | adjusted p-value | z score | combined score |
| regulation of protein targeting (GO:1903533) | 0.005936 | 0.03332 | -4.26 | 21.86 |
| ribosomal small subunit export from nucleus (GO:0000056) | 0.005936 | 0.03332 | -4.17 | 21.37 |
| mitochondrial electron transport, NADH to ubiquinone (GO:0006120) | 0.0007188 | 0.02581 | -2.74 | 19.8 |
| mitochondrial ATP synthesis coupled proton transport (GO:0042776) | 0.01854 | 0.04191 | -4.96 | 19.77 |
| mitochondrial ATP synthesis coupled electron transport (GO:0042775) | 0.00004997 | 0.003801 | -1.9 | 18.84 |
| folic acid-containing compound biosynthetic process (GO:0009396) | 0.005936 | 0.03332 | -3.45 | 17.69 |
| oxygen transport (GO:0015671) | 0.006781 | 0.03332 | -3.32 | 16.57 |
| positive regulation of autophagy of mitochondrion (GO:1903599) | 0.006781 | 0.03332 | -3 | 15 |
| ribosomal subunit export from nucleus (GO:0000054) | 0.011 | 0.04056 | -3.08 | 13.89 |
| pteridine-containing compound biosynthetic process (GO:0042559) | 0.006781 | 0.03332 | -2.68 | 13.41 |
| **Erythroid cluster \| CDK1i upregulated genes \| GO Biological Process** |  |  |  |  |
| Name | p-value | adjusted p-value | z score | combined score |
| negative regulation of membrane protein ectodomain proteolysis (GO:0051045) | 0.003196 | 0.04232 | -2.66 | 15.26 |
| negative regulation of reproductive process (GO:2000242) | 0.004392 | 0.04232 | -2.54 | 13.81 |
| sequestering of actin monomers (GO:0042989) | 0.004392 | 0.04232 | -2.43 | 13.17 |
| regulation of integrin-mediated signaling pathway (GO:2001044) | 0.004791 | 0.04232 | -2.3 | 12.27 |
| regulation of trophoblast cell migration (GO:1901163) | 0.004791 | 0.04232 | -2.24 | 11.96 |
| negative regulation of DNA replication (GO:0008156) | 0.006781 | 0.04303 | -2.2 | 10.96 |
| positive regulation of viral entry into host cell (GO:0046598) | 0.003595 | 0.04232 | -1.85 | 10.44 |
| transition metal ion homeostasis (GO:0055076) | 0.009958 | 0.04303 | -2.23 | 10.26 |
| cellular iron ion homeostasis (GO:0006879) | 0.02258 | 0.0497 | -2.61 | 9.91 |
| negative regulation of actin filament polymerization (GO:0030837) | 0.009165 | 0.04303 | -1.83 | 8.57 |

**Table 8.** GO terms for CDKli differentially expressed genes in the mesenchymal cluster.

| Mesenchymal cluster \| CDK1i downregulated genes \| GO Biological Process |  |  |  |  |
| --- | --- | --- | --- | --- |
| Name | p-value | adjusted p-value | z score | combined score |
| ribosomal small subunit export from nucleus (GO:0000056) | 0.005587 | 0.0459 | -4.17 | 21.63 |
| mitochondrial electron transport, NADH to ubiquinone (GO:0006120) | 0.0006352 | 0.0305 | -2.74 | 20.14 |
| mitochondrial ATP synthesis coupled proton transport (GO:0042776) | 0.01746 | 0.05732 | -4.96 | 20.08 |
| mitochondrial ATP synthesis coupled electron transport (GO:0042775) | 0.00004128 | 0.004197 | -1.9 | 19.21 |
| bleb assembly (GO:0032060) | 0.006383 | 0.0459 | -3 | 15.15 |
| ribosomal subunit export from nucleus (GO:0000054) | 0.01035 | 0.05579 | -3.08 | 14.08 |
| respiratory electron transport chain (GO:0022904) | 0.00005559 | 0.004197 | -1.36 | 13.32 |
| genetic imprinting (GO:0071514) | 0.007178 | 0.04927 | -2.56 | 12.64 |
| regulation of glycogen (starch) synthase activity (GO:2000465) | 0.006383 | 0.0459 | -2.43 | 12.27 |
| protein targeting to ER (GO:0045047) | 0.002727 | 0.03956 | -2.08 | 12.26 |

## REFERENCES

1. Tavian, M., Coulombel, L., Luton, D., Clemente, H.S., Dieterlen-Lièvre, F., and Péault, B. (1996). Aorta-associated CD34+ hematopoietic cells in the early human embryo. Blood 87, 67–72.

2. Medvinsky, A., and Dzierzak, E. (1996). Definitive hematopoiesis is autonomously initiated by the AGM region. Cell 86, 897–906.

3. Oberlin, E., Tavian, M., Blazsek, I., and Peault, B. (2002). Blood-forming potential of vascular endothelium in the human embryo. Development 129, 4147–4157.

4. Yokomizo, T., and Dzierzak, E. (2010). Three-dimensional cartography of hematopoietic clusters in the vasculature of whole mouse embryos. Development 137, 3651–3661.

5. Ivanovs, A., Rybtsov, S., Welch, L., Anderson, R.A., Turner, M.L., and Medvinsky, A. (2011). Highly potent human hematopoietic stem cells first emerge in the intraembryonic aorta-gonadmesonephros region. J. Exp. Med. 208, 2417–2427.

6. North, T., Gu, T.L., Stacy, T., Wang, Q., Howard, L., Binder, M., Marín-Padilla, M., and Speck, N.A. (1999). Cbfa2 is required for the formation of intra-aortic hematopoietic clusters. Development 126, 2563–2575.

7. Lancrin, C., Mazan, M., Stefanska, M., Patel, R., Lichtinger, M., Costa, G., Vargel, O., Wilson, N.K., Möröy, T., Bonifer, C., et al. (2012). GFI1 and GFI1B control the loss of endothelial identity of hemogenic endothelium during hematopoietic commitment. Blood 120, 314–322.

8. Wilson, N.K., Foster, S.D., Wang, X., Knezevic, K., Schütte, J., Kaimakis, P., Chilarska, P.M., Kinston, S., Ouwehand, W.H., Dzierzak, E., et al. (2010). Combinatorial Transcriptional Control In Blood Stem/Progenitor Cells: Genome-wide Analysis of Ten Major Transcriptional Regulators. Cell Stem Cell 7, 532–544.

9. Lichtinger, M., Ingram, R., Hannah, R., Müller, D., Clarke, D., Assi, S.A., Lie-A-Ling, M., Noailles, L., Vijayabaskar, M.S., Wu, M., et al. (2012). RUNX1 reshapes the epigenetic landscape at the onset of haematopoiesis. EMBO J. 31, 4318–4333.

10. Ottersbach, K., and Dzierzak, E. (2005). The murine placenta contains hematopoietic stem cells within the vascular labyrinth region. Dev. Cell 8, 377–387.

11. Robin, C., Bollerot, K., Mendes, S., Haak, E., Crisan, M., Cerisoli, F., Lauw, I., Kaimakis, P., Jorna, R., Vermeulen, M., et al. (2009). Human Placenta Is a Potent Hematopoietic Niche Containing Hematopoietic Stem and Progenitor Cells throughout Development. Cell Stem Cell 5, 385–395.

12. Li, Z., Lan, Y., He, W., Chen, D., Wang, J., Zhou, F., Wang, Y., Sun, H., Chen, X., Xu, C., et al. (2012). Mouse embryonic head as a site for hematopoietic stem cell development. Cell Stem Cell 11, 663–675.

13. Ivanovs, A., Rybtsov, S., Ng, E.S., Stanley, E.G., Elefanty, A.G., and Medvinsky, A. (2017). Human haematopoietic stem cell development: from the embryo to the dish. Development 144, 2323–2337.

14. Kumaravelu, P., Hook, L., Morrison, A.M., Ure, J., Zhao, S., Zuyev, S., Ansell, J., and Medvinsky, A. (2002). Quantitative developmental anatomy of definitive haematopoietic stem cells/longterm repopulating units (HSC/RUs): role of the aorta-gonad-mesonephros (AGM) region and the yolk sac in colonisation of the mouse embryonic liver. Development 129, 4891–4899.

15. Kaufman, D.S. (2009). Toward clinical therapies using hematopoietic cells derived from human pluripotent stem cells. Blood 114, 3513–3523.

16. Niwa, A., Heike, T., Umeda, K., Oshima, K., Kato, I., Sakai, H., Suemori, H., Nakahata, T., and Saito, M.K. (2011). A novel Serum-Free monolayer culture for orderly hematopoietic differentiation of human pluripotent cells via mesodermal progenitors. PLoS One 6.

17. Kennedy, M., Awong, G., Sturgeon, C.M., Ditadi, A., LaMotte-Mohs, Ross Zuniga-Pflucker, J.C., and Keller, G. (2012). T Lymphocyte Potential Marks the Emergence of Definitive Hematopoietic Progenitors in Human Pluripotent Stem Cell Differentiation Cultures. Cell Rep. 2, 1722–1735.

18. Slukvin, I.I. (2013). Hematopoietic specification from human pluripotent stem cells: current advances and challenges toward de novo generation of hematopoietic stem cells. Blood 122, 4035–4046.

19. Ramos-Mejía a, V., Navarro-Montero, O., Ayll??n, V., Bueno, C., Romero, T., Real, P.J., and Menendez, P. (2014). HOXA9 promotes hematopoietic commitment of human embryonic stem cells. Blood 124, 3065–3075.

20. Lis, R., Karrasch, C.C., Poulos, M.G., Kunar, B., Redmond, D., Duran, J.G.B., Badwe, C.R., Schachterle, W., Ginsberg, M., Xiang, J., et al. (2017). Conversion of adult endothelium to immunocompetent haematopoietic stem cells. Nature 545, 439–445.

21. Sugimura, R., Jha, D.K., Han, A., Soria-Valles, C., da Rocha, E.L., Lu, Y.-F., Goettel, J.A., Serrao, E., Rowe, R.G., Malleshaiah, M., et al. (2017). Haematopoietic stem and progenitor cells from human pluripotent stem cells. Nature 545, 432–438.

22. Sturgeon, C.M., Ditadi, A., Awong, G., Kennedy, M., and Keller, G. (2014). Wnt signaling controls the specification of definitive and primitive hematopoiesis from human pluripotent stem cells. Nat. Biotechnol. 32, 554–561.

23. Ditadi, A., Sturgeon, C.M., Tober, J., Awong, G., Kennedy, M., Yzaguirre, A.D., Azzola, L., Ng, E.S., Stanley, E.G., French, D.L., et al. (2015). Human definitive haemogenic endothelium and arterial vascular endothelium represent distinct lineages. Nat. Cell Biol. 17, 580–591.

24. Boiers, C., Carrelha, J., Lutteropp, M., Luc, S., Green, J.C.A., Azzoni, E., Woll, P.S., Mead, A.J., Hultquist, A., Swiers, G., et al. (2013). Lymphomyeloid contribution of an immune-restricted progenitor emerging prior to definitive hematopoietic stem cells. Cell Stem Cell 13, 535–548.

25. McGrath, K.E., Frame, J.M., Fegan, K.H., Bowen, J.R., Conway, S.J., Catherman, S.C., Kingsley, P.D., Koniski, A.D., and Palis, J. (2015). Distinct Sources of Hematopoietic Progenitors Emerge before HSCs and Provide Functional Blood Cells in the Mammalian Embryo. Cell Rep. 11, 1892–1904.

26. Butler, A., Hoffman, P., Smibert, P., Papalexi, E., and Satija, R. (2018). Integrating single-cell transcriptomic data across different conditions, technologies, and species. Nat. Biotechnol. 36, 411–420.

27. ten Dijke, P., Egorova, A.D., Goumans, M.-J.T.H., Poelmann, R.E., and Hierck, B.P. (2012). TGF-β Signaling in Endothelial-to-Mesenchymal Transition: The Role of Shear Stress and Primary Cilia. Sci. Signal. 5, pt2–pt2.

28. Chen, P.Y., Qin, L., Baeyens, N., Li, G., Afolabi, T., Budatha, M., Tellides, G., Schwartz, M.A., and Simons, M. (2015). Endothelial-to-mesenchymal transition drives atherosclerosis progression. J. Clin. Invest. 125, 4514–4528.

29. Good, R.B., Gilbane, A.J., Trinder, S.L., Denton, C.P., Coghlan, G., Abraham, D.J., and Holmes, A.M. (2015). Endothelial to Mesenchymal Transition Contributes to Endothelial Dysfunction in Pulmonary Arterial Hypertension. Am. J. Pathol. 185, 1850–1858.

30. Zhong, A., Mirzaei, Z., and Simmons, C.A. (2018). The Roles of Matrix Stiffness and ß-Catenin Signaling in Endothelial-to-Mesenchymal Transition of Aortic Valve Endothelial Cells. Cardiovasc. Eng. Technol. 9, 158–167.

31. Man, S., Sanchez Duffhues, G., ten Dijke, P., and Baker, D. (2018). The therapeutic potential of targeting the endothelial-to-mesenchymal transition. Angiogenesis.

32. Ivanovs, A., Rybtsov, S., Anderson, R.A., and Medvinsky, A. (2014a). CD43 but Not CD41 Marks the First Hematopoietic Stem Cells in the Human Embryo. Blood 124.

33. Ivanovs, A., Rybtsov, S., Anderson, R.A., Turner, M.L., and Medvinsky, A. (2014b). Identification of the niche and phenotype of the first human hematopoietic stem cells. Stem Cell Reports 2, 449–456.

34. Watt, S.M., Butler, L.H., Tavian, M., Bühring, H.J., Rappold, I., Simmons, P.J., Zannettino, a C., Buck, D., Fuchs, a, Doyonnas, R., et al. (2000). Functionally defined CD164 epitopes are expressed on CD34(+) cells throughout ontogeny but display distinct distribution patterns in adult hematopoietic and nonhematopoietic tissues. Blood 95, 3113–3124.

35. Ohata, S., Nawa, M., Kasama, T., Yamasaki, T., Sawanobori, K., Hata, S., Nakamura, T., Asaoka, Y., Watanabe, T., Okamoto, H., et al. (2009). Hematopoiesis-dependent expression of CD44 in murine hepatic progenitor cells. Biochem. Biophys. Res. Commun. 379, 817–823.

36. Cao, H., Heazlewood, S.Y., Williams, B., Cardozo, D., Nigro, J., Oteiza, A., and Nilsson, S.K. (2016). The role of CD44 in fetal and adult hematopoietic stem cell regulation. Haematologica 101, 26–37.

37. Kozar, K., Ciemerych, M.A., Rebel, V.I., Shigematsu, H., Zagozdzon, A., Sicinska, E., Geng, Y., Yu, Q., Bhattacharya, S., Bronson, R.T., et al. (2004). Mouse Development and Cell Proliferation in the Absence of D-Cyclins. Cell 118, 477–491.

38. Sakaue-Sawano, A., Kurokawa, H., Morimura, T., Hanyu, A., Hama, H., Osawa, H., Kashiwagi, S., Fukami, K., Miyata, T., Miyoshi, H., et al. (2008). Visualizing spatiotemporal dynamics of multicellular cell-cycle progression. Cell 132, 487–498.

39. Pauklin, S., and Vallier, L. (2013). The cell cycle state of stem cells determines cell fate propensity. Cell 155, 135–147.

40. Blajeski, A.L., Phan, V.A., Kottke, T.J., and Kaufmann, S.H. (2002). G(1) and G(2) cell-cycle arrest following microtubule depolymerization in human breast cancer cells. J. Clin. Invest. 110, 91–99.

41. Yiangou, L., Grandy, R.A., Morell, C.M., Tomaz, R.A., Osnato, A., Kadiwala, J., Muraro, D., Garcia-Bernardo, J., Nakanoh, S., Bernard, W.G., et al. (2019). Method to Synchronize Cell Cycle of Human Pluripotent Stem Cells without Affecting Their Fundamental Characteristics. Stem Cell Reports 12, 165–179.

42. Zhang, J., Li, H., Yabut, O., Fitzpatrick, H., D’Arcangelo, G., and Herrup, K. (2010). Cdk5 suppresses the neuronal cell cycle by disrupting the E2F1-DP1 complex. J. Neurosci. 30, 5219–5228.

43. Su, S.C., and Tsai, L.-H. (2011). Cyclin-Dependent Kinases in Brain Development and Disease. Annu. Rev. Cell Dev. Biol. 27, 465–491.

44. Hydbring, P., Malumbres, M., and Sicinski, P. (2016). Non-canonical functions of cell cycle cyclins and cyclin-dependent kinases. Nat. Rev. Mol. Cell Biol. 17, 280–292.

45. Krentz, N.A.J., van Hoof, D., Li, Z., Watanabe, A., Tang, M., Nian, C., German, M.S., and Lynn, F.C. (2017). Phosphorylation of NEUROG3 Links Endocrine Differentiation to the Cell Cycle in Pancreatic Progenitors. Dev. Cell 41, 129–142.e6.

46. Meijer, L., Borgne, A., Mulner, O., Chong, J.P., Blow, J.J., Inagaki, N., Inagaki, M., Delcros, J.G., and Moulinoux, J.P. (1997). Biochemical and cellular effects of roscovitine, a potent and selective inhibitor of the cyclin-dependent kinases cdc2, cdk2 and cdk5. Eur. J. Biochem. 243, 527–536.

47. Knockaert, M., Gray, N., Damiens, E., Chang, Y.-T., Grellier, P., Grant, K., Fergusson, D., Mottram, J., Soete, M., Dubremetz, J.-F., et al. (2000). Intracellular targets of cyclin-dependent kinase inhibitors: identification by affinity chromatography using immobilised inhibitors. Chem. Biol. 7, 411–422.

48. Vassilev, L.T. (2006). Cell Cycle Synchronization at the G2/M Phase Border by Reversible Inhibition of CDK1. Cell Cycle 5, 2555–2556.

49. McInnes, C. (2008). Progress in the evaluation of CDK inhibitors as anti-tumor agents. Drug Discov. Today 13, 875–881.

50. Finn, R.S., Dering, J., Conklin, D., Kalous, O., Cohen, D.J., Desai, A.J., Ginther, C., Atefi, M., Chen, I., Fowst, C., et al. (2009). PD 0332991, a selective cyclin D kinase 4/6 inhibitor, preferentially inhibits proliferation of luminal estrogen receptor-positive human breast cancer cell lines in vitro. Breast Cancer Res. 11, R77.

51. Rocca, A., Farolfi, A., Bravaccini, S., Schirone, A., and Amadori, D. (2014). Palbociclib (PD 0332991): targeting the cell cycle machinery in breast cancer. Expert Opin. Pharmacother. 15, 407–420.

52. Vella, S., Tavanti, E., Hattinger, C.M., Fanelli, M., Versteeg, R., Koster, J., Picci, P., and Serra, M. (2016). Targeting CDKs with Roscovitine Increases Sensitivity to DNA Damaging Drugs of Human Osteosarcoma Cells. PLoS One 11, e0166233.

53. Prevo, R., Pirovano, G., Puliyadi, R., Herbert, K.J., Rodriguez-Berriguete, G., O’Docherty, A., Greaves, W., McKenna, W.G., and Higgins, G.S. (2018). CDK1 inhibition sensitizes normal cells to DNA damage in a cell cycle dependent manner. Cell Cycle 1–11

54. Li, X., Rong, Y., Zhang, M., Wang, X.L., LeMaire, S.A., Coselli, J.S., Zhang, Y., and Shen, Y.H. (2009). Up-regulation of thioredoxin interacting protein (Txnip) by p38 MAPK and FOXO1 contributes to the impaired thioredoxin activity and increased ROS in glucose-treated endothelial cells. Biochem. Biophys. Res. Commun. 381, 660–665.

55. Liu, J.-Y., Yao, J., Li, X.-M., Song, Y.-C., Wang, X.-Q., Li, Y.-J., Yan, B., and Jiang, Q. (2014). Pathogenic role of lncRNA-MALAT1 in endothelial cell dysfunction in diabetes mellitus. Cell Death Dis. 5, e1506.

56. Fuschi, P., Carrara, M., Voellenkle, C., Garcia-Manteiga, J.M., Righini, P., Maimone, B., Sangalli, E., Villa, F., Specchia, C., Picozza, M., et al. (2017). Central role of the p53 pathway in the noncoding-RNA response to oxidative stress. Aging (Albany. NY). 9, 2559–2586.

57. Tian, D., Dong, J., Jin, S., Teng, X., and Wu, Y. (2017). Endogenous hydrogen sulfide-mediated MAPK inhibition preserves endothelial function through TXNIP signaling. Free Radic. Biol. Med. 110, 291–299.

58. Malumbres, M., Sotillo, R., Santamarıa, D., Galan, J., Cerezo, A., Ortega, S., Dubus, P., and Barbacid, M. (2004). Mammalian Cells Cycle without the D-Type Cyclin-Dependent Kinases Cdk4 and Cdk6. Cell 118, 493–504.

59. Batsivari, A., Rybtsov, S., Souilhol, C., Binagui-Casas, A., Hills, D., Zhao, S., Travers, P., and Medvinsky, A. (2017). Understanding Hematopoietic Stem Cell Development through Functional Correlation of Their Proliferative Status with the Intra-aortic Cluster Architecture. Stem Cell Reports 8, 1549–1562.

60. Fujimoto, T., Anderson, K., Jacobsen, S.E.W., Nishikawa, S., and Nerlov, C. (2007). Cdk6 blocks myeloid differentiation by interfering with Runx1 DNA binding and Runx1-C/EBPα interaction. EMBO J. 26, 2361–2370.

61. Taoudi, S. (2005). Progressive divergence of definitive haematopoietic stem cells from the endothelial compartment does not depend on contact with the foetal liver. Development 132, 4179–4191.

62. Yusa, K., Rashid, S.T., Strick-Marchand, H., Varela, I., Liu, P.-Q., Paschon, D.E., Miranda, E., Ordonez, A., Hannan, N.R.F., Rouhani, F.J., et al. (2011). Targeted gene correction of α1-antitrypsin deficiency in induced pluripotent stem cells. Nature 478, 391–394.

63. Chen, G., Gulbranson, D.R., Hou, Z., Bolin, J.M., Ruotti, V., Probasco, M.D., Smuga-Otto, K., Howden, S.E., Diol, N.R., Propson, N.E., et al. (2011b). Chemically defined conditions for 168 human iPSC derivation and culture. Nat. Methods 8, 424–429.

64. Zeng, Y., He, J., Bai, Z., Li, Z., Gong, Y., Liu, C., Ni, Y., Du, J., Ma, C., Bian, L., Lan, Y., & Liu, B. (2019). Tracing the first hematopoietic stem cell generation in human embryo by single-cell RNA sequencing. Cell Research, 29(11), 881–894.

65. Stuart, T., Butler, A., Hoffman, P., Hafemeister, C., Papalexi, E., Mauck, W.M., Hao, Y., Stoeckius, M., Smibert, P., and Satija, R. (2019). Comprehensive Integration of Single-Cell Data. Cell 177, 1888–1902.e21.

66. Bertero, A., Pawlowski, M., Ortmann, D., Snijders, K., Yiangou, L., De Brito, M.C., Brown, S., Bernard, W.G., Cooper, J.D., Giacomelli, E., et al. (2016). Optimized inducible shRNA and CRISPR/Cas9 platforms for in vitro studies of human development using hPSCs. Dev. 143, 4405–4418.

67. Wei, Y., Chen, Y.H., Li, L.Y., Lang, J., Yeh, S.P., Shi, B., Yang, C.C., Yang, J.Y., Lin, C.Y., Lai, C.C., et al. (2011). CDK1-dependent phosphorylation of EZH2 suppresses methylation of H3K27 and promotes osteogenic differentiation of human mesenchymal stem cells. Nat. Cell Biol. 13, 87–94.

68. Yiangou, L., Grandy, R.A., Osnato, A., Ortmann, D., Sinha, S., and Vallier, L. (2019). Cell cycle regulators control mesoderm specification in human pluripotent stem cells. J. Biol. Chem. 294, 17903–17914.

