## Supplementary figures and images for "Analysis of Endothelial-to-Haematopoietic Transition at the Single Cell Level identifies Cell Cycle Regulation as a Driver of Differentiation"

### Supplementary Fig 1

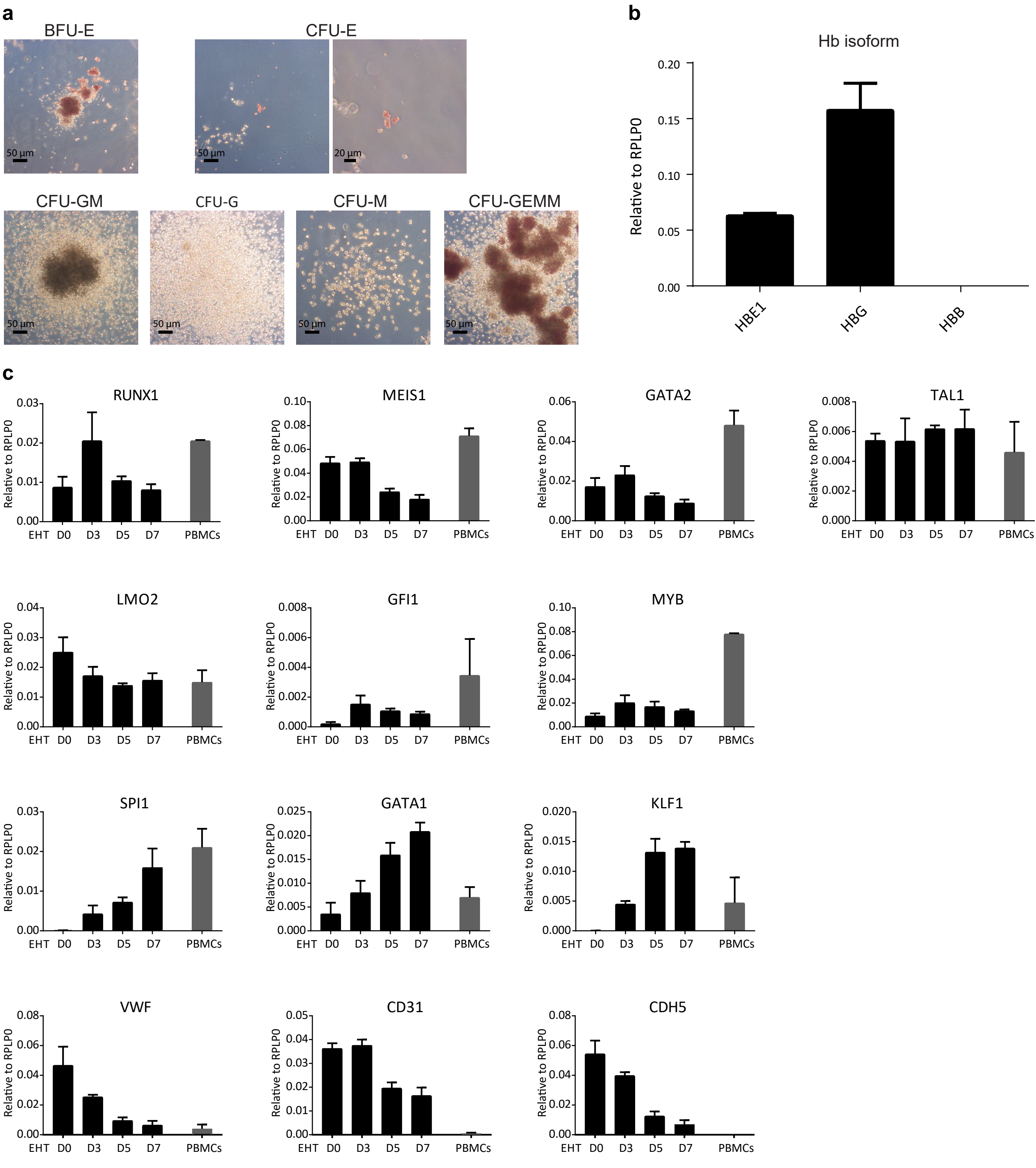

### Supplementary Fig 2

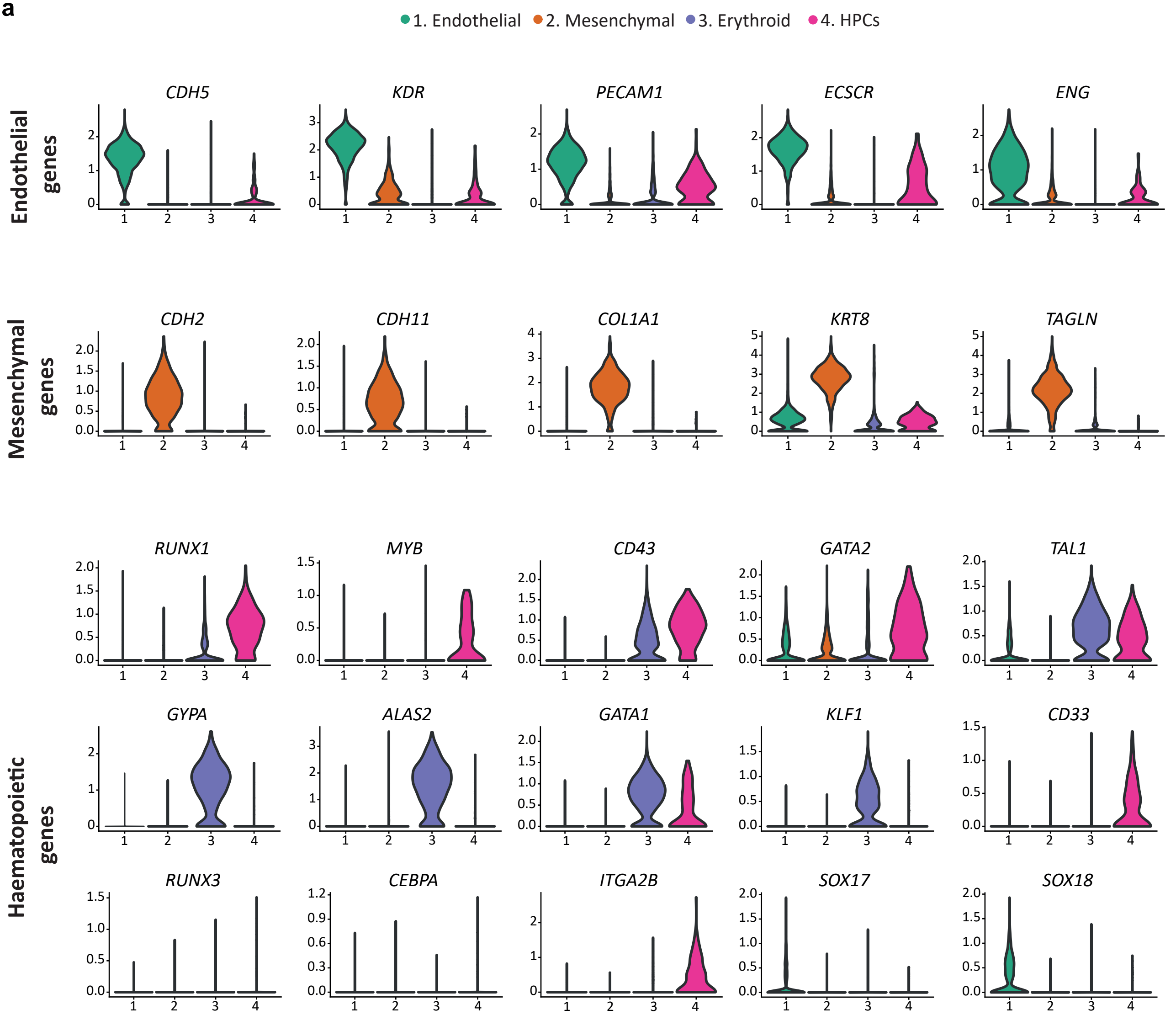

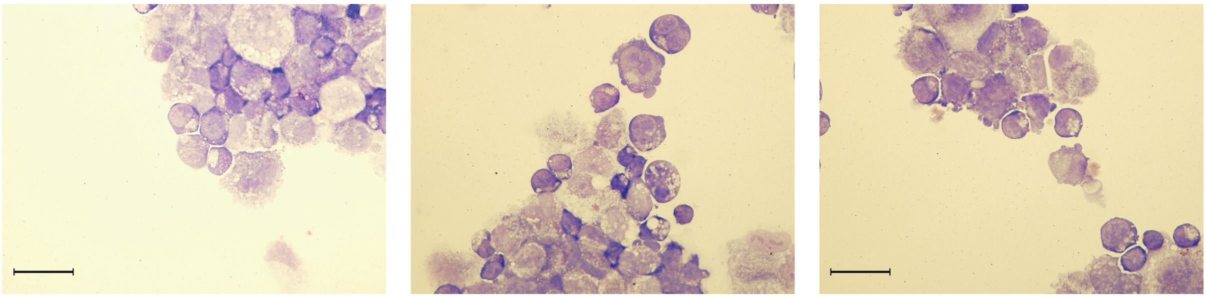

### Supplementary Fig 3

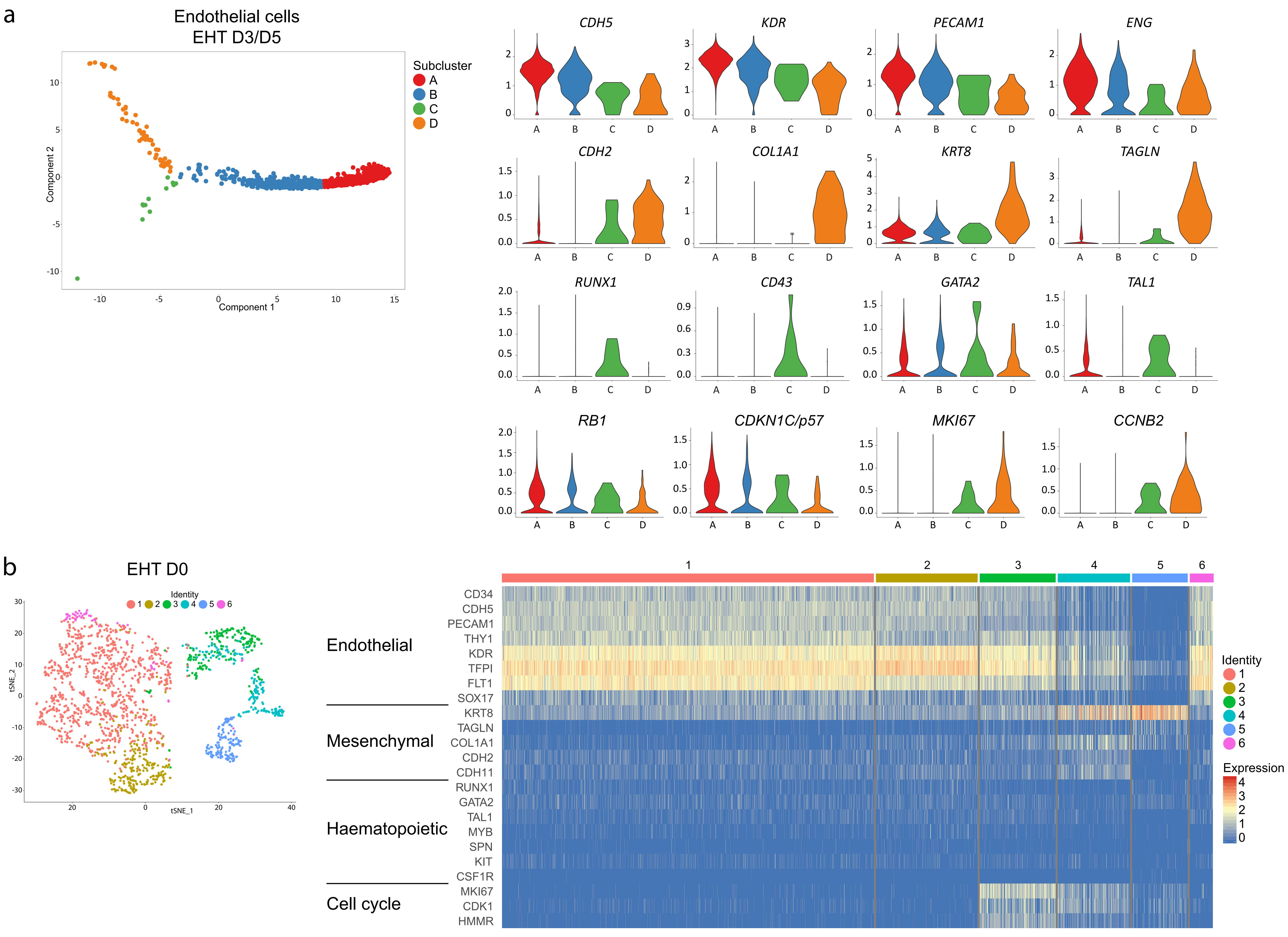

### Supplementary Fig 4

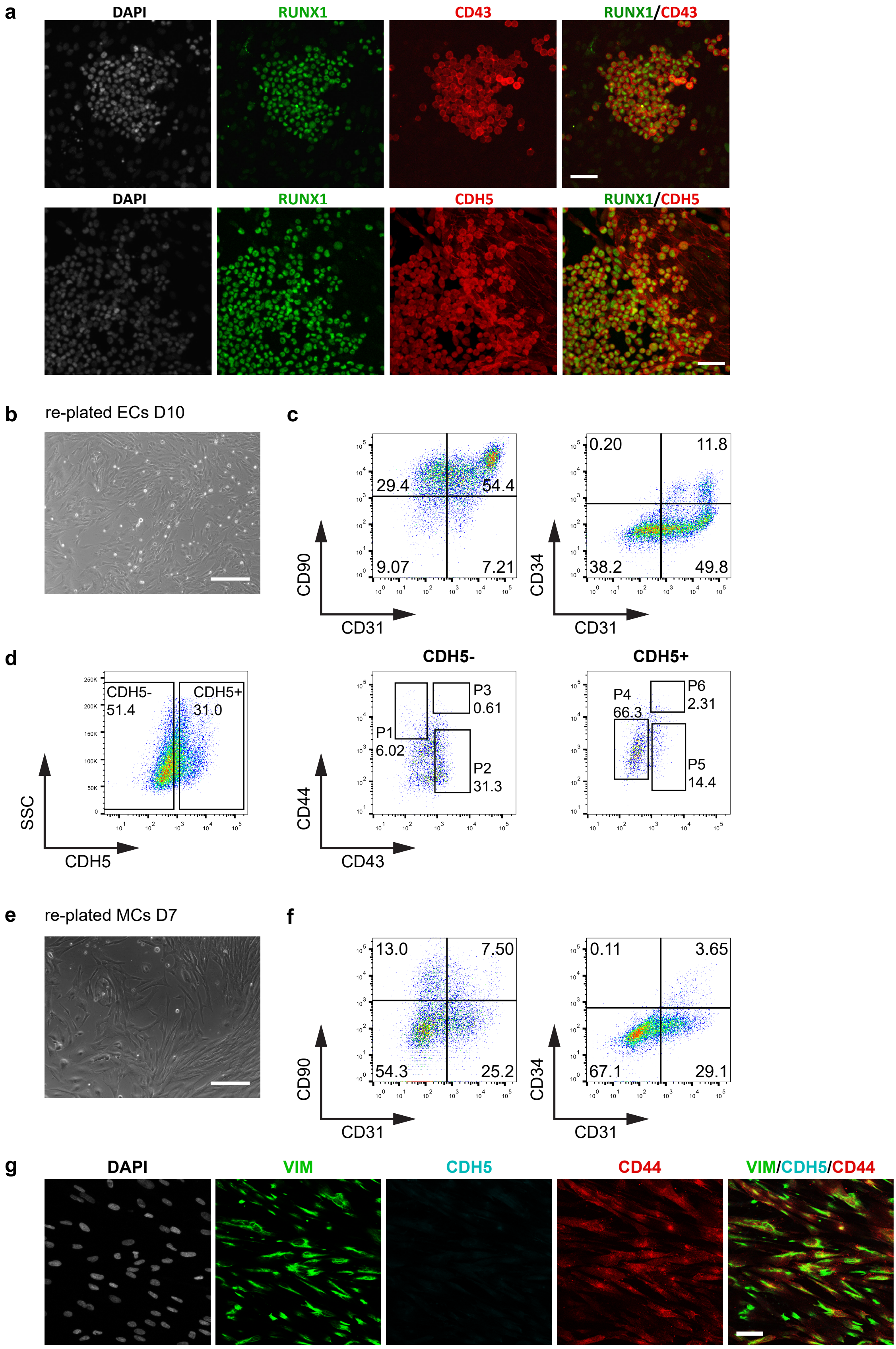

### Supplementary Fig 5

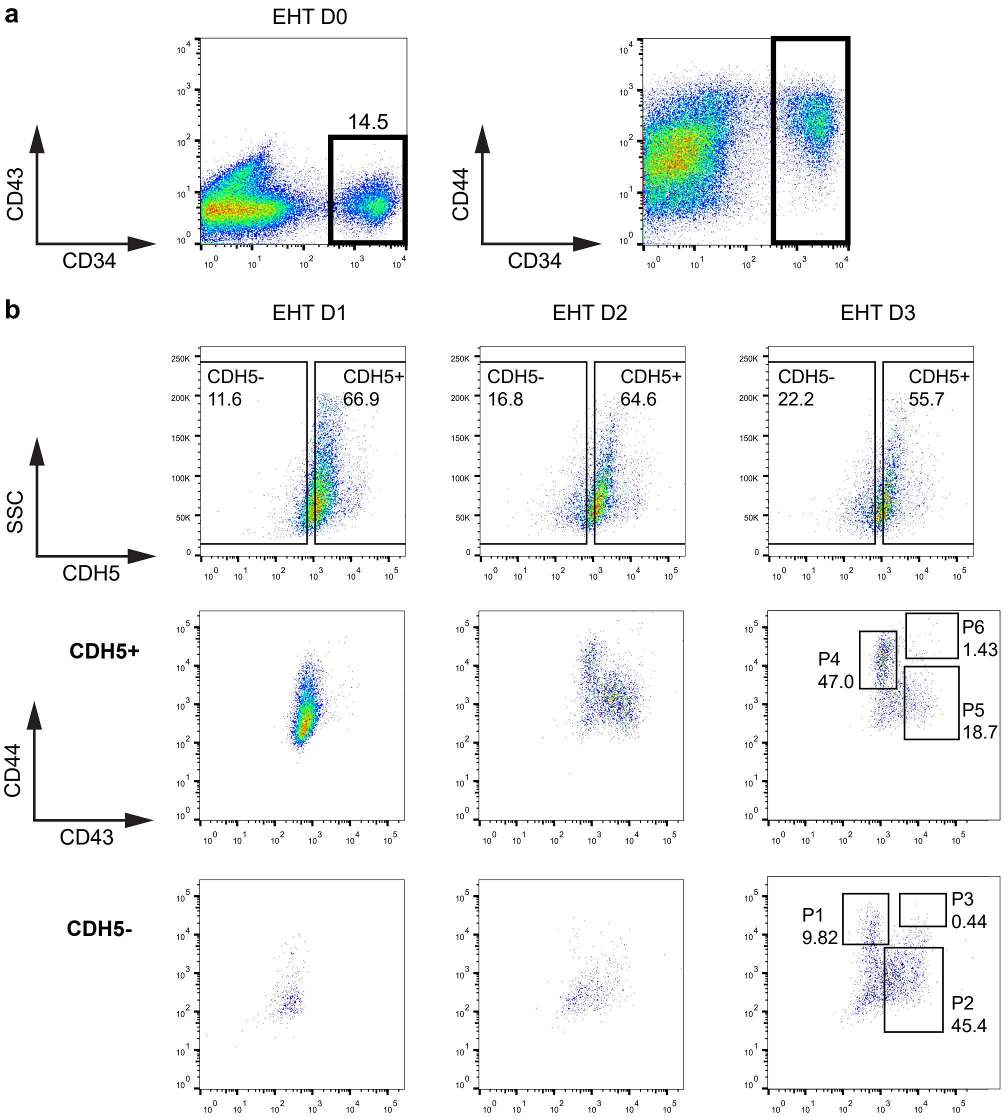

### Supplementary Fig 6

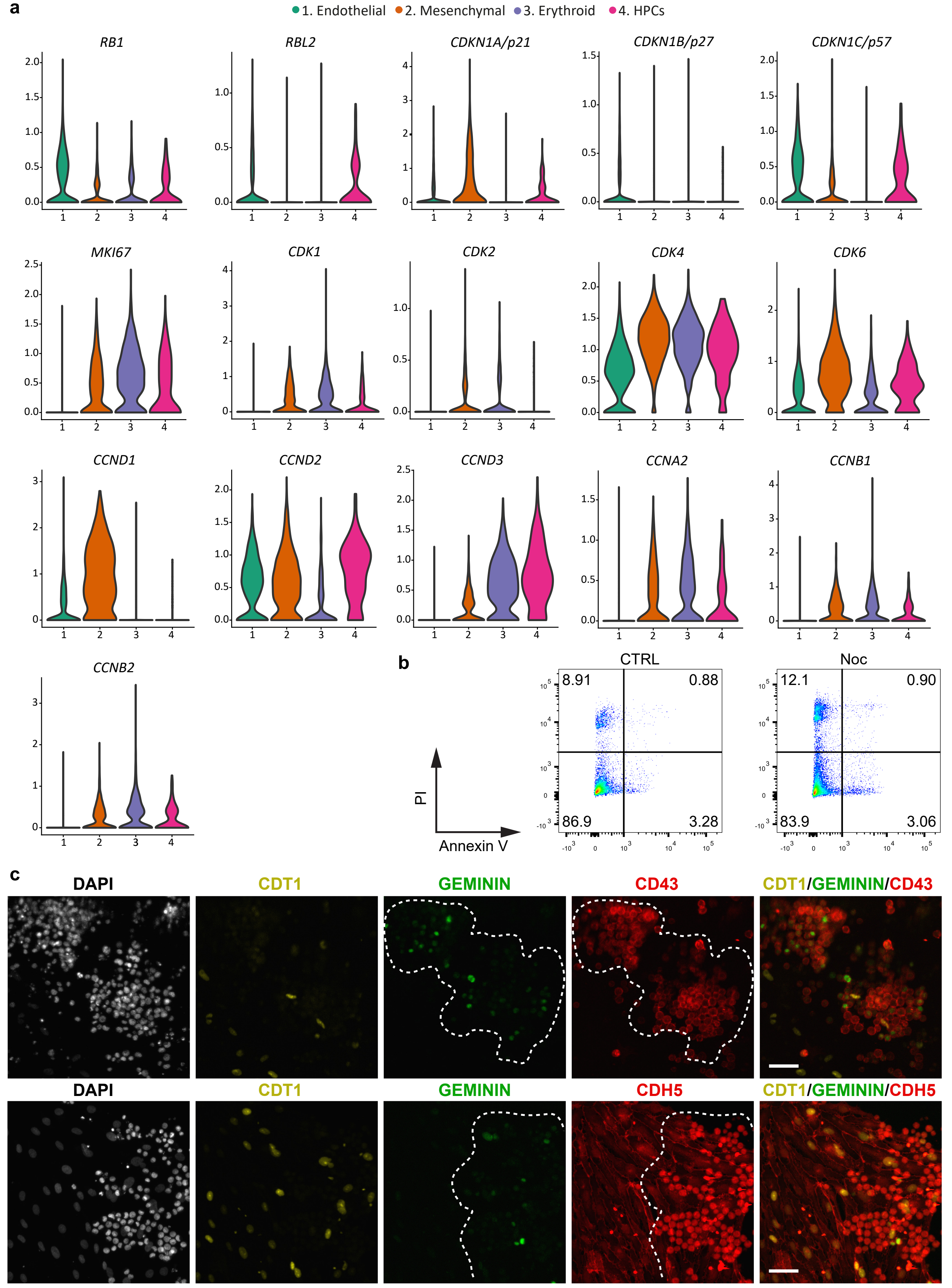

### Supplementary Fig 7

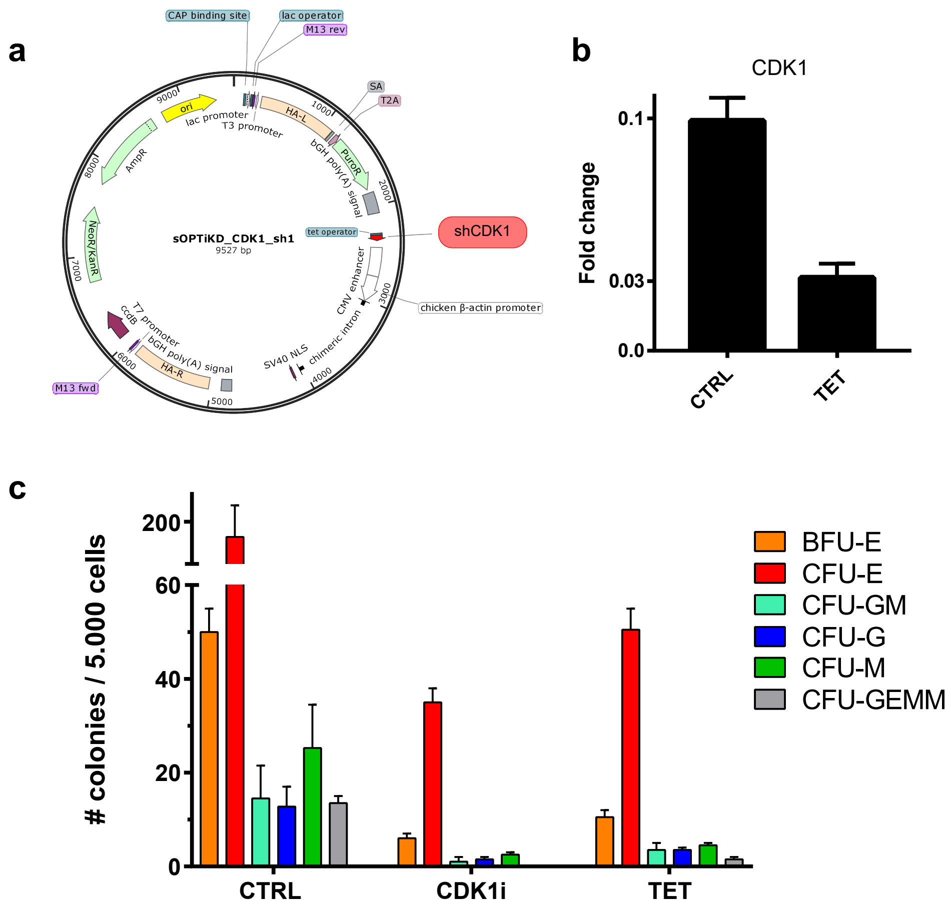

### Supplementary Fig 8

CTRL *vs* CDK4/6i

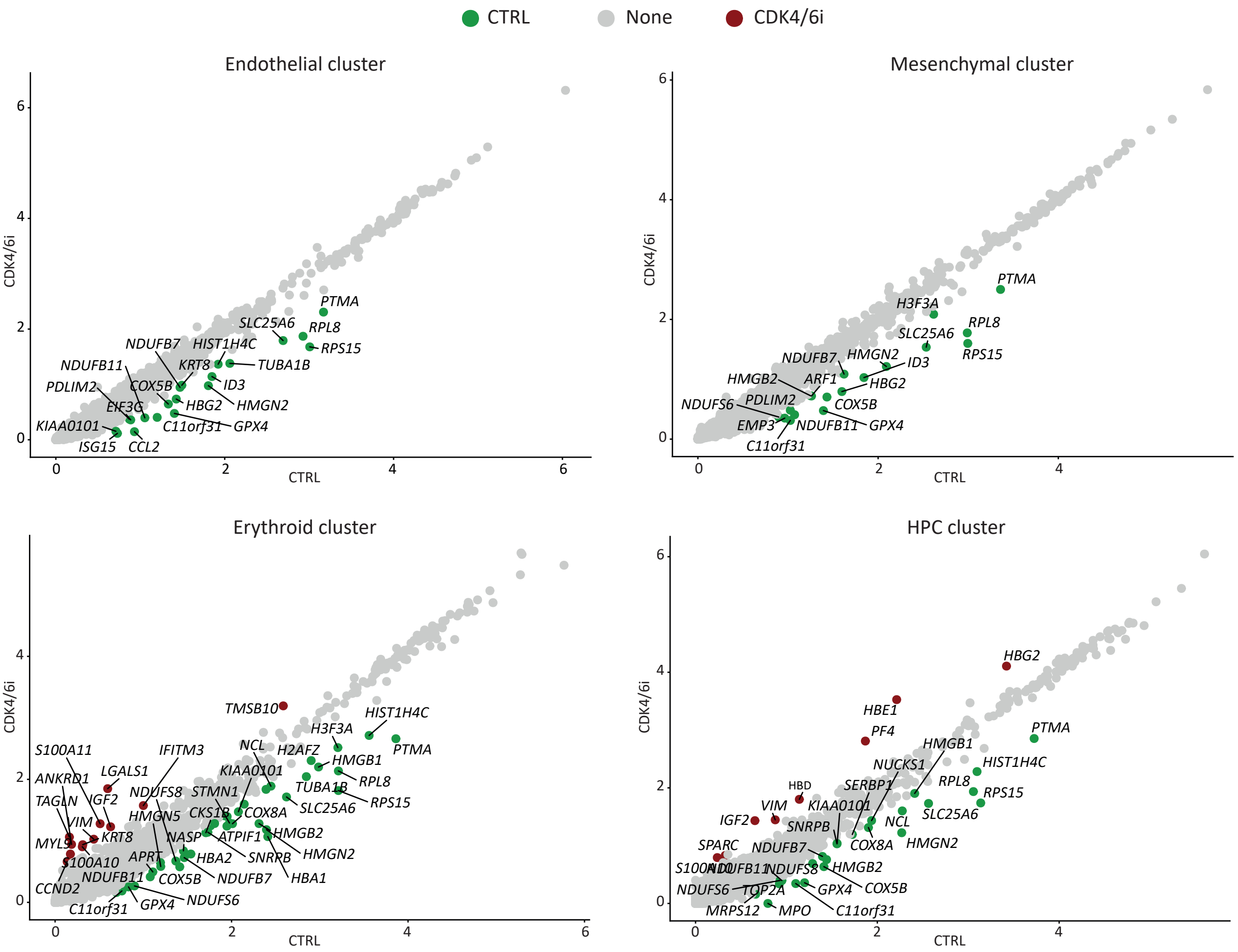


CTRL *vs* CDK1i

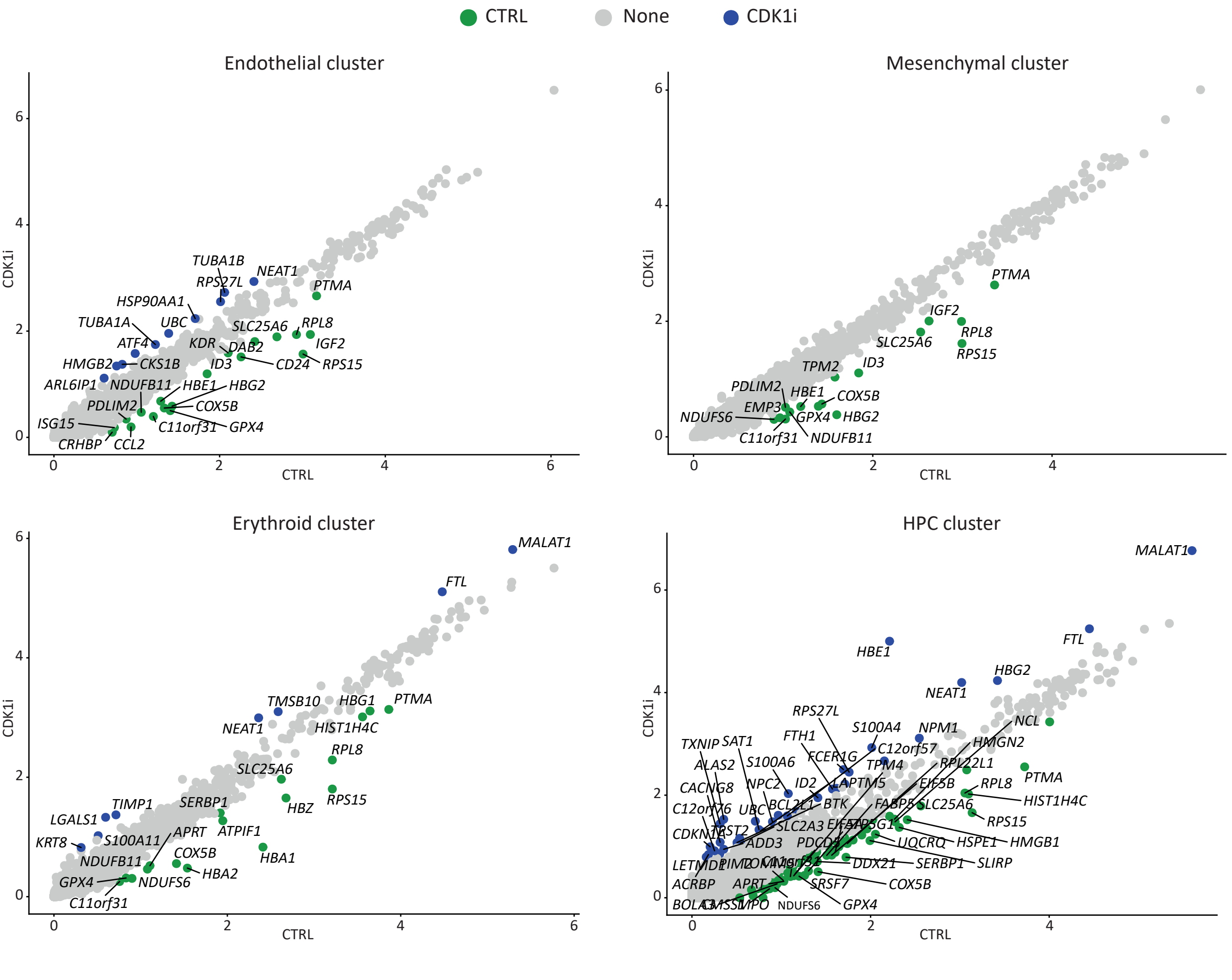

### Supplementary Fig 9

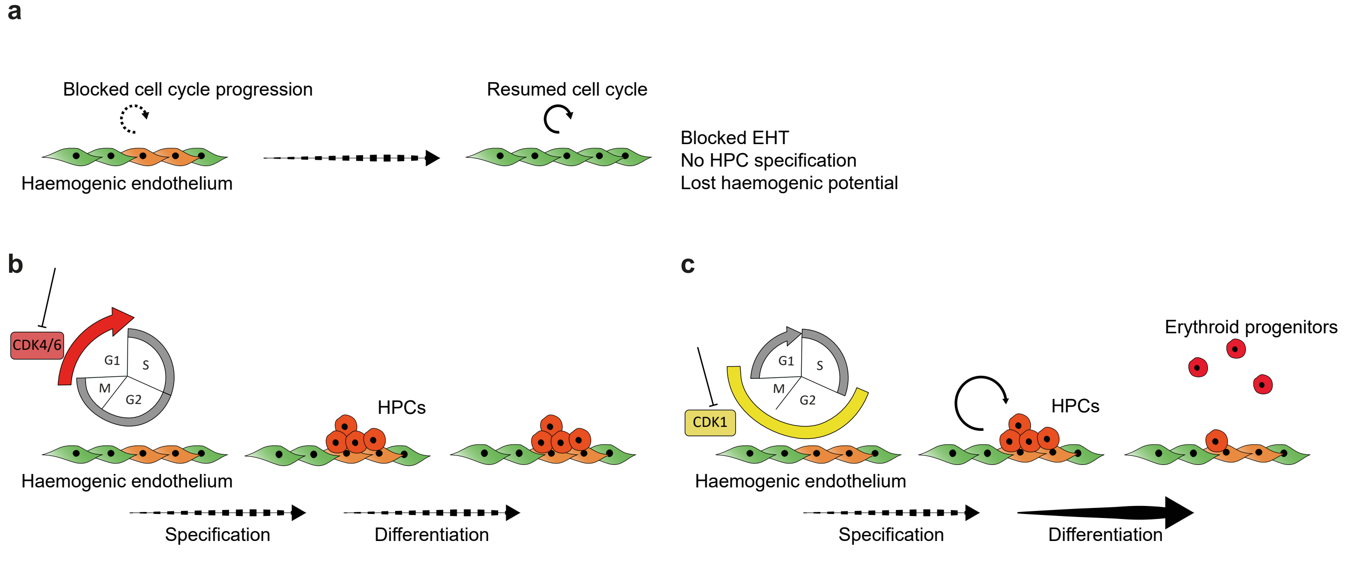

### Supplementary Fig 10

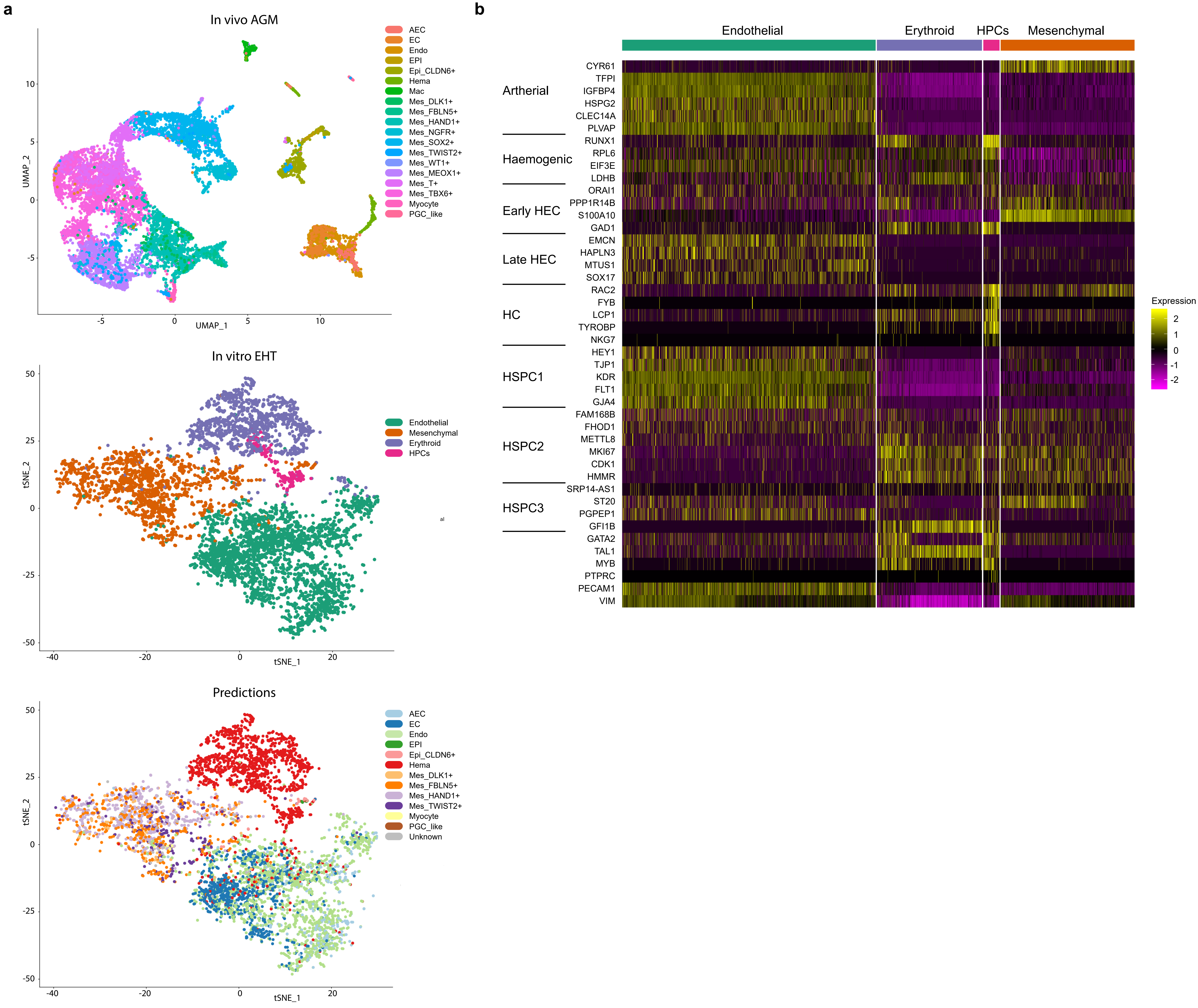
